# Non-additive interactions between multiple mutualists and host plant genotype simultaneously promote increased plant growth and pathogen defense

**DOI:** 10.1101/2025.02.26.640321

**Authors:** Amanda H. Rawstern, Lucas J. Carbajal, Tyler J. Slade, Michelle E. Afkhami

## Abstract

Understanding the impact of microbial interactions on plants is critical for maintaining healthy native ecosystems and sustainable agricultural practices. Despite the reality that genetically distinct plants host multiple microbes-of-large effect in the field, it remains unclear the extent to which host genotypes modulate non-additive microbial interactions and how these interactions differ between benign/pathogenic environments. Our study fills this gap by performing a large-scale manipulative microbiome experiment across 7 genotypes of the model legume *Medicago truncatula*. We combine plant performance metrics, survival analyses, predictive modeling, RNA extractions, and targeted gene expression to assess how host genotype and microbes non-additively interact to shape plant growth and disease ecology. Our results reveal three important findings: (1) host genotypes with high tolerance to pathogens benefit more from multiple mutualist interactions than susceptible genotypes, (2) mutualists confer the same non-additive plant performance benefits in both benign and pathogenic environments, and (3) the quality of the symbiotic relationship with mutualists is a strong predictor of host survival against pathogenic disease. By applying these findings towards developing crops that promote synergistic microbial interactions, yields and pathogen defense could be simultaneously increased while reducing the need for toxic fertilizers and pesticides.

## Introduction

Plant pathogens are a devastating threat to both native and agro-ecosystems and have become increasingly prominent due to the exacerbated climate pressures and globalization of commerce and travel within the Anthropocene (Anderson et al. 2004, Raffa et al. 2023, Singh et al. 2023, Strange and Scott 2005). Over the last decade, pathogenic fungi have nearly extirpated whole families of once common forest trees including Laurels (Choudhury et al. 2021), Beeches (Boyd et al. 2013), and Elms (Boyd et al. 2013); and pathogens alone have caused up to 40% yield reduction and over $200 billion USD losses in major agricultural crops (Savary et al. 2019).

Routine management practices include the application of chemicals, such as bactericides and fungicides (Sundin et al. 2016, Zubrod et al. 2019). These practices are not sustainable as they have not only led to unintended consequences, such as polluting waterways with toxic chemicals (De Souza et al. 2020), but are also costly given that reliance on chemicals requires investing in a perpetual arms race with pathogens that often become resistant (Cycoń, Mrozik, and Piotrowska-Seget 2019, Sundin et al. 2016). Alternative strategies proposed include the manipulation of microbiomes and host-associated microbial mutualists to promote plant health and prime plant immune responses (Poudel et al. 2016, Trivedi et al. 2020), as well as the targeted genetic modification of plants to increase their resistance to pathogens (Dong and Ronald 2019, Esse, Reuber, and Does 2020).

Beneficial microbes are important hidden players of disease ecology as they can alter all three cornerstones of the classic disease triangle model: 1) virulence of the pathogen, 2) susceptibility of the host, and 3) conduciveness of the environment (Leveau 2024, McNew 1960). First, beneficial microbes can decrease pathogen virulence by secreting antimicrobial compounds that inhibit pathogen survival (Chen et al. 2009), outcompete pathogens for limited resources (Kinkel et al. 2012), and physically block access to the host (Sassone-Corsi 2015, Vigo, Norman, and Hooker 2000). Second, mutualists can make hosts more robust against pathogen invasion by increasing host nutrition, growth, and overall health (Dordas 2008, Karagiannidis, Bletsos, and Stavropoulos 2002, Rahman et al. 2018), and by stimulating biochemical changes associated with host defense mechanisms that prime the host immune system (Pozo et al. 2002). Third, beneficial microbes can also change the environment making it less conducive to pathogen invasion, by modifying abiotic soil conditions like pH and water retention (Philippot et al. 2024) or by recruiting a community of microbiota “unfavorable” to the pathogen (Hoge 2000). This research has been an integral foundation for identifying major pathways through which microbes *individually* mitigate pathogen stress. However, in the complex reality of plant disease ecology, plants are impacted by a variety of key symbionts that can switch from being beneficial to detrimental depending on their non-additive interactions with other microbes (Afkhami, Rudgers, and Stachowicz 2014, Afkhami et al. 2020).

Non-additive effects of microbes on the performance of hosts are ubiquitous (Almeida, Tran, and Afkhami 2024, Baichman-Kass, Song, and Friedman 2023) and cannot be predicted based on single microbe studies alone. Non-additive effects can range from synergism (i.e. joint effect on host performance is greater than additive summation of individual microbial effects) to antagonism (i.e. joint effect on host performance is less than additive summation). A recent meta-analysis suggests that plants generally benefit from co-inoculation of multiple mutualists, but that plant biomass can range from being twice as small to over three times as large as uninoculated plants depending on factors including plant age and phenology (Kasanke et al. 2024). Another plant attribute that impacts multiple mutualist interactions is genotype. Distinct plant genotypes can interact with microbes differentially due to variation in root exudates (Sasse, Martinoia, and Northern, 2018) and plant cell wall surface receptors (Bulgarelli et al. 2013).

Despite this, most studies utilize only one genotype in their experimental design and rarely quantify population-level variation (Franklin, Hockey, and Maherali et al. 2020), which is thought to be one of the major contributing factors for the low efficacy of inoculation benefits among diverse populations in the field (Kaminsky et al. 2019, Maged, Abdul, and Heribert 2020). Therefore, it would be of value to understand the extent to which host plant genotypes modulate non-additive microbial interactions and how these interactions impact pathogen susceptibility or resistance.

Legumes, which are considered to be one of the most important families of plants due to their diverse roles in agriculture and ecosystem health and resilience, not only experience substantial negative effects of microbial pathogens but also participate in some of the most important and widespread microbial mutualisms on Earth (Lodwig et al. 2003, Libault et al. 2009, Tian, Kah, and Kariman 2019). Legumes include popular foods such as the common bean, chickpeas, soybeans, peanuts, and lentils that are crucial protein sources sustaining human populations around the world as well as forage, like alfalfa and clover, that is vital to both livestock and wild grazers. The lignocellulosic biomass of discarded legume pods and hulls have also been shown to be a promising source of biofuel production (Ankita et al. 2023). Further, legumes form important symbioses with arbuscular mycorrhizal fungi (AMF), which often helps hosts increase nutrient and water uptake (Bisht, Sharma, and Garg 2024), and with bacterial rhizobia, which fix atmospheric nitrogen into ammonium providing ∼65% of the biosphere’s available nitrogen (Lodwig et al. 2003). Foundational insights garnered from research on the model legume *Medicago truncatula* (Barrel medic) have been successfully applied to the improvement and management of agricultural legume crops (Gou et al. 2018, Kumar et al. 2015, Roorkiwal et al. 2020, Young and Udvardi 2009). Previously, work from our lab has found that multiple mutualists act synergistically in benign environments to increase the strength of selection on plant architectural traits, which emphasizes that multiple mutualists not only immediately benefit host performance but also shape legume evolutionary trajectories (Afkhami, Friesen, and Stinchcombe 2021). However, it remains unclear how multiple mutualists and plant genotype interact to impact host performance and whether these interactions confer similar host performance outcomes in a pathogenic environment.

In the current study, we aim to address these gaps by performing a large-scale manipulative experiment of common multiple microbial mutualists *(Rhizophagus irregularis* and *Ensifer melioti*) and a pervasive soil-borne fungal pathogen (*Fusarium oxysporum*) across 7 genotypes of the model legume *Medicago truncatula* which cluster into groups that range from highly tolerant to highly susceptible to pathogenic infection. We use plant performance metrics (i.e. photosynthetic leaf count, branch count, and plant height), survival analyses, predictive modeling, RNA extractions, and targeted gene expression to assess how multiple mutualists interact with host tolerance genotype to shape the severity of pathogenic disease. Specifically, we tested: (1) Do multiple mutualist non-additive effects on plant performance differ across plant tolerance genotype and are these effects consistent between benign and pathogenic environments?, (2) How does the interaction between plant tolerance genotype and symbiont inoculation impact plant survival against pathogenic attack and can aboveground plant performance metrics prior to pathogen invasion be used to predict plant survival?, and (3) How does the presence of a pathogen impact mutualist abundances and symbioses quality within plant root tissues? Taken together, our results show non-additivity of multiple mutualists is prevalent, host genotypes with high tolerance to pathogens benefit more from multiple mutualist interactions than susceptible genotypes, mutualists confer the same host performance outcomes in both benign and pathogenic environments, and host survival of a fungal pathogen attack is dependent on the quality of the symbiotic relationship with multiple mutualists. Overall, this work elucidates how complex interactions between nitrogen-fixing bacteria, arbuscular mycorrhizal fungi, and plant tolerance genotype shape legume disease ecology.

## Materials and Methods

### Experimental design

To determine the role that host genotype plays in the frequency of non-additive interactions, we selected 7 unique *Medicago truncatula* genotypes that cluster into three groups: high, medium, and low tolerance to fungal pathogens (Table S1; Fig. 1A). These genotypes have been previously classified into these groupings based on either their differing levels of antifungal saponin production (Lei et al. 2019) and/or their fungal infection survival rates (Rispail and Rubiales 2014). To assess the impact of multiple mutualist inoculation on legume plant performance metrics in benign and pathogenic environments, we planted 960 plants in a completely randomized block design (Fig. 1B) with a 2x2x2 factorial manipulation of the presence/absence of AMF (*Rhizophagus irregularis* DAOM197198) (M±), rhizobia (*Ensifer meiloti* Rm1021) (R±), and fusarium pathogen (*Fusarium oxysporum f.sp. medicaginis* 5190a)(P±).

**Fig. 1.**
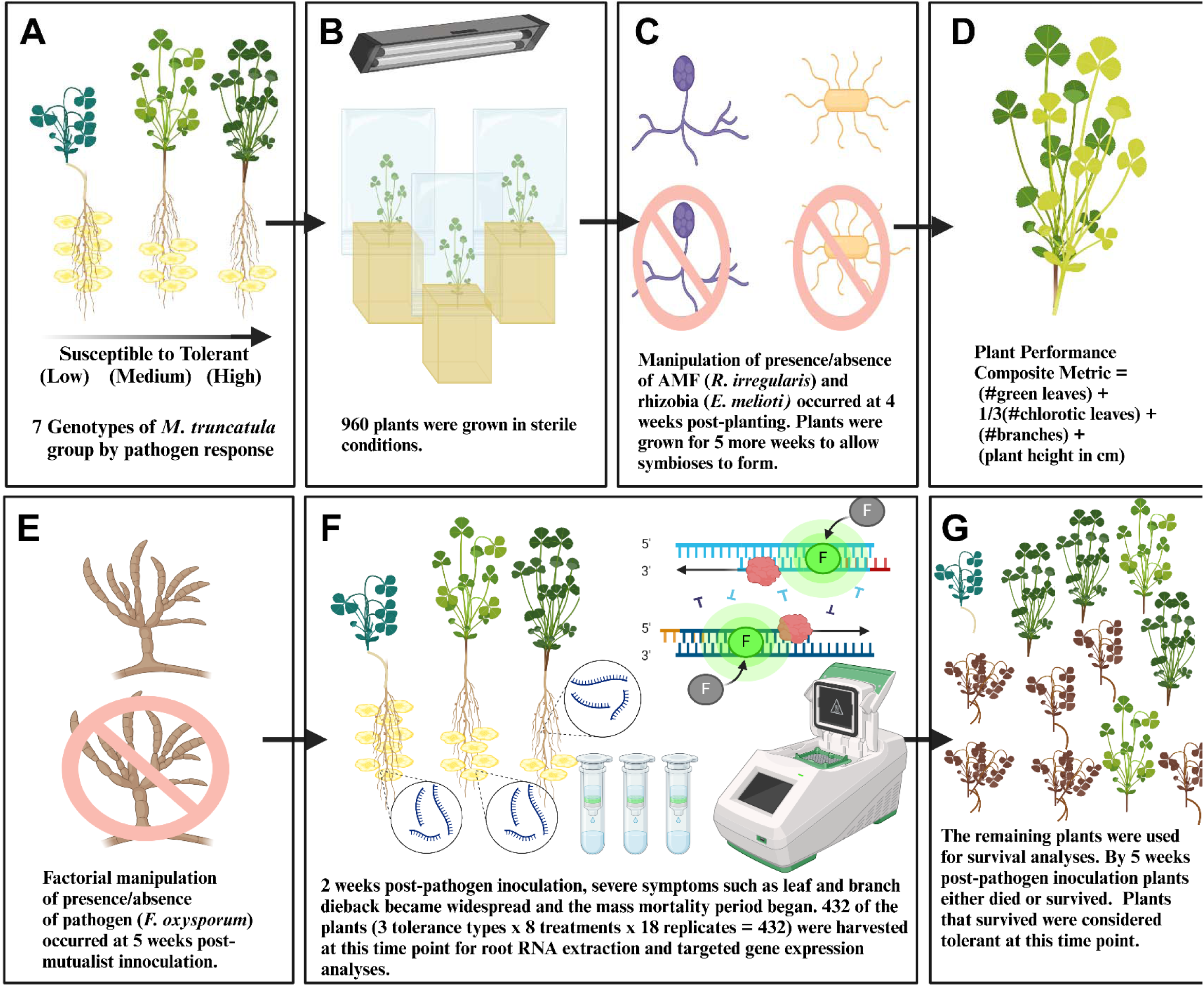
Experimental design overview. **(A)** We selected 7 *M. truncatula* genotypes that group into low tolerance (susceptible), medium tolerance, and high tolerance (resistant) to general fungal pathogen invasion. **(B)** Our experiment used 960 plants in a 2x2x2 factorial manipulation of the presence/absence of arbuscular mycorrhizal fungi (AMF) (*R. irregularis*; M±), rhizobia (*E. meiloti*; R±), and fusarium pathogen (*F. oxysporum*; P±). **(C)** After inoculation with AMF and rhizobia, plants were grown for 5 weeks to allow symbioses to occur, and the first data set was collected. **(D)** Data collected was a composite metric of plant performance by summing center-scaled normalized metrics of photosynthetic leaves, number of branches, and plant height. **(E)** Plants were inoculated with *F. oxysporum*, a globally important soil-borne pathogen that infects plant root water conducting tissues. **(F)** 2 weeks post-pathogen inoculation the second data set is collected. This is the time point where severe symptoms such as leaf and branch dieback became widespread. 432 plants (3 tolerance types x 8 treatments x 18 replicates = 432 plants) were harvested for RNA extraction and targeted gene expression analyses. **(G)** The remaining plants were used for survival analyses. By 5 weeks post-pathogen inoculation plants have either died or survived and survivors reach maturation for seeding. The last data set on plant performance was collected at this time point. Figure created using BioRender.com.

### Preparation of germinants and microbial inoculants

Seeds were obtained from the National Center for Genome Resources (Santa Fe, NM, USA) and the USDA-ARS Legume Germplasm Repository (Pullman, WA, USA). Seeds were mechanically scarified, surface disinfected, and sterilely germinated following standard protocol (Garcia, Barker, and Journet 2006), and then aseptically planted in magenta boxes containing a 1:1 volume ratio of sand to perlite to prevent crushing of the roots (Barker et al. 2006). To create a closed system, we sterilized the sand-perlite mixture in the magenta boxes in an autoclave at 121°C three times prior to planting and attached a sterile bag over the top of each magenta box (Afkhami and Stinchcombe 2016) (Fig. 1B). Plant growth conditions were 25°C with a light cycle of 16 hours light/8hours dark. All plants were watered once per week with 5 mL of filter-sterilized low nitrogen/phosphorus fertilizer with remaining macronutrients optimized for *Medicago truncatula* (De Bang et al. 2017).

The rhizobia were grown to log phase (i.e. exponential growth phase with highest concentration of viable cells) in liquid media (Journet et al. 2006) and diluted to a concentration of ∼10^6^ cells/mL (OD_600_ = 0.1) in sterile water (Simonsen and Stinchcombe 2014). AMF spores were commercially purchased through Premier Tech (Rivière-du-Loup, QC, CA) and diluted to a concentration of ∼300 spores/mL in sterile water (Afkhami and Stinchcombe 2016) using a hemocytometer. Each R+ plant received 1 mL of rhizobia inoculant (∼10^6^ cells) while each M+ plant received 1 mL (∼300 spores) of AMF (Fig. 1C). All plants then received additional sterile water (either 3, 4, or 5 mL depending on treatment group) so that every plant received 5 mL of total liquid. Inoculation of mutualists occurred at 4 weeks after initial planting to allow time for secondary root elongation.

*Fusarium oxysporum f.sp. medicaginis* pathogen was obtained from Dr. Deborah Samac of the USDA-ARS Plant Science Research Unit (St. Paul, MN, USA) under USDA-APHIS permit P526-21-05187. The pathogen was grown in liquid media until sporulation (Lichtenzveig et al. 2006) and diluted to a concentration of ∼300 spores/mL in sterile water using a hemocytometer. Each P+ plant received 1 mL (∼300 spores) of fusarium pathogen. All plants then received either 4 or 5 mL of sterile water so that every plant received 5 mL of total liquid (Fig. 1E). Inoculation of the pathogen occurred at 5 weeks post-inoculation with the mutualists to give the plants time to form both rhizobia nodules (Journet et al. 2006) and mycorrhizal arbuscules (Chabaud et al. 2006). *Fusarium oxysporum* was chosen as the pathogen model because it is an ubiquitous and globally important soil-borne pathogen with over 100 host-specific strains and has been detrimental to numerous agroecosystems by infecting the xylem water-conducting tissues of roots (Gordon 2017).

### Non-additivity of plant performance and survival analyses

To understand the interaction between plant genotype and multiple mutualists on aboveground plant performance, we calculated a composite performance metric by summing center-scaled normalized metrics (Subedi et al. 2022) of the number of photosynthetic leaves, number of branches, and plant height (in cm) as a proxy for aboveground growth. The number of photosynthetic leaves was calculated using the following equation: (number of green leaves) + 1/3(number of chlorotic leaves) based upon previous work in *Medicago truncatula* that determined green leaves had around three times the photosynthetic capacity as chlorotic leaves (Chuanen et al. 2011). Data collection of plant performance occurred at three critical timepoints: 5 weeks post-inoculation with the mutualists (time point by which symbiosis with both mutualists typically occurs; Journet et al. 2006, Chabaud et al. 2006; Fig. 1D), 2 weeks after fusarium pathogen inoculation (time point at which pathogen switches to being necrotrophic, severe symptoms such as leaf and branch dieback become widespread, and mass mortality begins; Rispail and Rubiales 2014; Fig. 1F), and 5 weeks after fusarium pathogen inoculation (time point at which plants have either died or survived and survivors reach maturation for seeding; Rispail and Rubiales 2014, Sultana et al. 2016; Fig. 1G).

For the non-additivity analyses, we confirmed that genotypes within each tolerance cluster performed the same (i.e. no significant interaction p > 0.05 of genotypes and treatment or main effect of genotype on the composite plant performance metric within the same tolerance cluster), supporting our clustering of the data into the three tolerance types. The composite plant performance metric for inoculated plants was normalized by calculating the percent change in the composite plant performance metric from the average of the controls uninoculated with mutualists (R-M-plants pre-pathogen/R-M-P+ plants post-pathogen). The mean for the additive expectation was determined by adding the mean for the rhizobia only group (R+M-plants pre-pathogen/R+M-P+ plants post-pathogen) to the mean of the AMF only group (R-M+ plants pre-pathogen/R-M+P+ post-pathogen). The standard error of the sum of two means was calculated using the basic statistical formula: 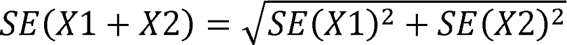 and the 95% CI was calculated as the mean ± z*SE, where a critical z-score of 1.96 was used.

Mutualists of plants inoculated with both types (R+M+ plants pre-pathogen/R+M+P+ post-pathogen) were considered synergistic if above the additive expectation CI, antagonistic if below the additive expectation CI, and additive if within the additive expectation CI. All statistical analyses were performed using R (v4.3.2; R Core Team 2023). To compare whether the frequency of non-additive interactions differs among plant genotype groups in a benign environment, we performed chi square tests of homogeneity between the high, medium, and low tolerance clusters from the data collected at the first critical timepoint (i.e. 5 weeks post-inoculation with mutualists). To determine whether the frequencies of non-additive plant performance responses change in a pathogenic environment, we performed Monte Carlo simulations (Waller et al. 2003) between the comparable pairs (i.e. high tolerance cluster in benign environment to high tolerance cluster in pathogenic environment, etc.) using the data collected at the second critical timepoint (i.e. 2 weeks post-inoculation with fusarium). To compare the change in plant growth through time repeated measure ANOVAs were used.

The survival rate for mutualist inoculated plants was normalized by calculating the percent change in the survival rate from the average of the controls uninoculated with mutualists (R-M-P+ plants) from data collected at the third critical time point (i.e. 5 weeks post-inoculation with fusarium). Survival to this timepoint was the focus of our analyses because this was the timepoint at which survivors are considered to have resisted the pathogen (Rispail and Rubiales 2014) and reach maturation for seed production (Sultana et al. 2016). Therefore, only plants that were able to survive to this stage were able to have a reproductive fitness beyond zero. To assess whether aboveground plant performance prior to pathogen inoculation could be used to predict plant survival after pathogen exposure, we performed a logistic regression and determined model performance using conventional k-fold cross-validation (Stone 1974). Our survival model performed with a k-fold cross-validation accuracy of 0.73, indicating the model has a good fit according to current machine learning modeling standards (Maciejauskaite and Miliauskaite 2024). Statistical differences in plant size and survival were determined using ANOVAs and chi-square proportion tests respectively.

### RNA root tissue collection and extraction

A subset of the plants (3 tolerance types x 8 treatments x 18 replicates = 432 plants) were harvested for RNA extraction at 2 weeks post-fusarium inoculation which is the period when severe disease symptoms occur but immediately before mass mortality sets in (Fig. 1F). Roots were flash frozen in dry ice and subsequently transferred to a -80°C freezer for storage. Full root systems were homogenized using a mortar and pestle in liquid nitrogen. Total RNA was extracted from each root sample using the Qiagen RNeasy PowerPlant Kit following the manufacturer’s protocol (Carlsbad, CA, USA). RNA was purified using magnetic bead cleaning and quantity was checked using a Qiagen Qubit 4 fluorometer. Negative controls showed that no cross-contamination occurred. DNA was depleted from the samples using Thermo Scientific DNase I (Waltham, MA, USA). cDNA was then generated using the Bio-Rad iScript cDNA Synthesis Kit (Hercules, CA, USA) utilizing the kit-provided random primers and 10 µL of RNA template. Before use, the cDNA was also purified using standard magnetic bead cleaning.

### Microbial abundance and targeted gene expression analyses

To evaluate how the presence of a pathogen impacts symbioses in plant root tissues, abundances for each microbe were measured by targeting common housekeeping genes (actin genes for eukaryotes and smc00128 gene for rhizobia) with previously validated primers specific to *Rhizophagus irregularis* (Tskuzki et al. 2016), *Fusarium oxyporum f.sp. medicaginis* (Williams et al. 2016), and *Ensifer melioti* (Krol and Becker 2004). Housekeeping genes are genes that are expressed at consistent transcription levels even under different environmental conditions and therefore the abundance of these transcripts strongly correlate with the abundance of active cells (Milanese et al. 2019). To quantify the number of active cells, standard dilution curves using a log10 scale were calculated for each microbe by extracting RNA and preparing cDNA from known masses of pure culture (Fig. S1), employing the same protocols as in the above section “RNA root tissue collection and extraction”. For quantification, qPCR was performed with Bio-Rad SsoAdvanced Universal SYBR Green Supermix using 10 µL supermix, 0.5 µL each (10x stock) of forward and reverse primers, 6.5 µL of nuclease-free water, and 2.5 µL of cDNA template. Reactions were performed in a Bio-Rad Cfx96 thermocycler with the conditions: 30s at 95°C for polymerase activation, 40 cycles of 15s at 95°C for denaturation and 45s at 60°C for annealing/extension/plate read, followed by standard melt curve analysis. The “predict” function in base R was used to extrapolate microbial abundances from the standard curves based on the sample’s Ct value.

To evaluate how the presence of a pathogen impacts host plant symbiosis quality with the mutualists, relative expression analyses were performed on three symbiotic marker genes of *Medicago truncatula* using previously validated primers (Breuillin-Sessoms et al. 2015, Liu et al. 2007). Upregulation of PT8 (AMF induced phosphorus transporter gene) and AMT 2;3 (AMF induced ammonium transporter gene) have been associated with increased phosphorus and nitrogen uptake in plant roots and shoots and are indicative of high-quality symbioses from beneficial AMF species (Breuillin-Sessoms et al. 2015, Cope et al. 2022). In addition, AMT 2;3 also plays an important role in regulating the lifespan of arbuscules by preventing premature arbuscule degeneration (Breuillin-Sessoms et al. 2015). TC100851 (symbiont upregulated multifunctional aquaporin gene) is one of the few membrane transporters upregulated in both nodulating and mycorrhizal roots (Benedito et al. 2010) and has also been associated with disease resistance against a bacterial pathogen (Liu et al. 2007). In general, multifunctional aquaporins have been found to increase water transport in plants, increase absorption of symbiont-conferred nutrients, and have an important role in preventing dehydration during pathogenic attack (Tyerman, Niemietz, and Bramley 2002, Wang et al. 2018). The relative expressions of these symbiotic marker genes were calculated using the delta Ct method (Fleige et al. 2006) with the *Medicago truncatula* actin housekeeping gene (Ivashuta et al. 2005) used as the internal control. These genes were measured on a log2 scale since every difference of -1Ct value represents a doubling of gene product (Liu et al. 2007). The same qPCR master mix and cycling conditions were used for these analyses as described for the microbial abundance genes above.

Full models evaluating factor impact on microbial abundance or symbiotic marker gene expression that were assessed contained the explanatory variables of ‘Microbial Treatment Group’ (microbe treatment added to plant), ‘Plant Tolerance’ (high, medium, low tolerance to fungal pathogens), ‘Pathogen Status’ (presence or absence of fusarium pathogen inoculation) and their interactions. All model explanatory variables were assessed for significance using type II ANOVAs which follows best practices for designs with unbalanced groups (Langsrud 2003). Comparisons in relative expression among significantly different groups were calculated using delta delta Ct (Livak and Schmittgen 2001).

## Results

### High tolerance plants have the highest rate of synergism while low tolerance plants have the highest rate of antagonism from interaction with multiple mutualists

We found that >77% of individuals responded non-additively with respect to plant performance when inoculated with multiple mutualists (5 weeks post-inoculation with mutualists; Fig. 2). We discovered that plant performance non-additivity response was significantly shaped by plant genotype, which clustered into high, medium, or low tolerance to fungal pathogens (chi-square test of homogeneity: X^2^ = 13.79, p = 0.008; Fig.2). Each tolerance cluster showed a different distribution of non-additivity responses with high and low tolerance clusters being distinct from one another (pairwise chi-square post-hoc test: X^2^ = 11.27, p = 0.003), while the medium group displayed intermediate characteristics (Fig. 2). This suggests that genotypes that fall within the same cluster have shared genetic characteristics that make them not only similar in their tolerance to fungal pathogens but also similar in their response to multiple mutualists. For the high tolerance cluster, the most common response was synergistic (51%, Fig. 2A) where plants with multiple mutualists performed ∼400-1000% better than controls and this response was significantly more common than additive (20%, p < 0.001, Fig. 2A) or antagonistic responses (29%, p = 0.01, Fig 2A). In the medium tolerance cluster, synergistic, additive, and antagonistic responses were all equally distributed (∼33% each, p = 0.85, Fig. 2B). Within the low tolerance cluster, the most common response was antagonistic (53%, Fig. 2C, Fig. S2) where plants with multiple symbionts performed up to ∼400% worse than controls, and this response was significantly more common than additive (20%, p < 0.001, Fig. 2C) or synergistic (27%, p = 0.002, Fig. 2C). Interestingly, these findings indicate that genotypes with high tolerance to fungal pathogens have the highest benefits/lowest costs to interacting with multiple mutualists, while genotypes with low tolerance experience the lowest benefits/highest costs when interacting with multiple mutualists, and medium tolerance genotypes show intermediate benefits/costs.

**Fig. 2.**
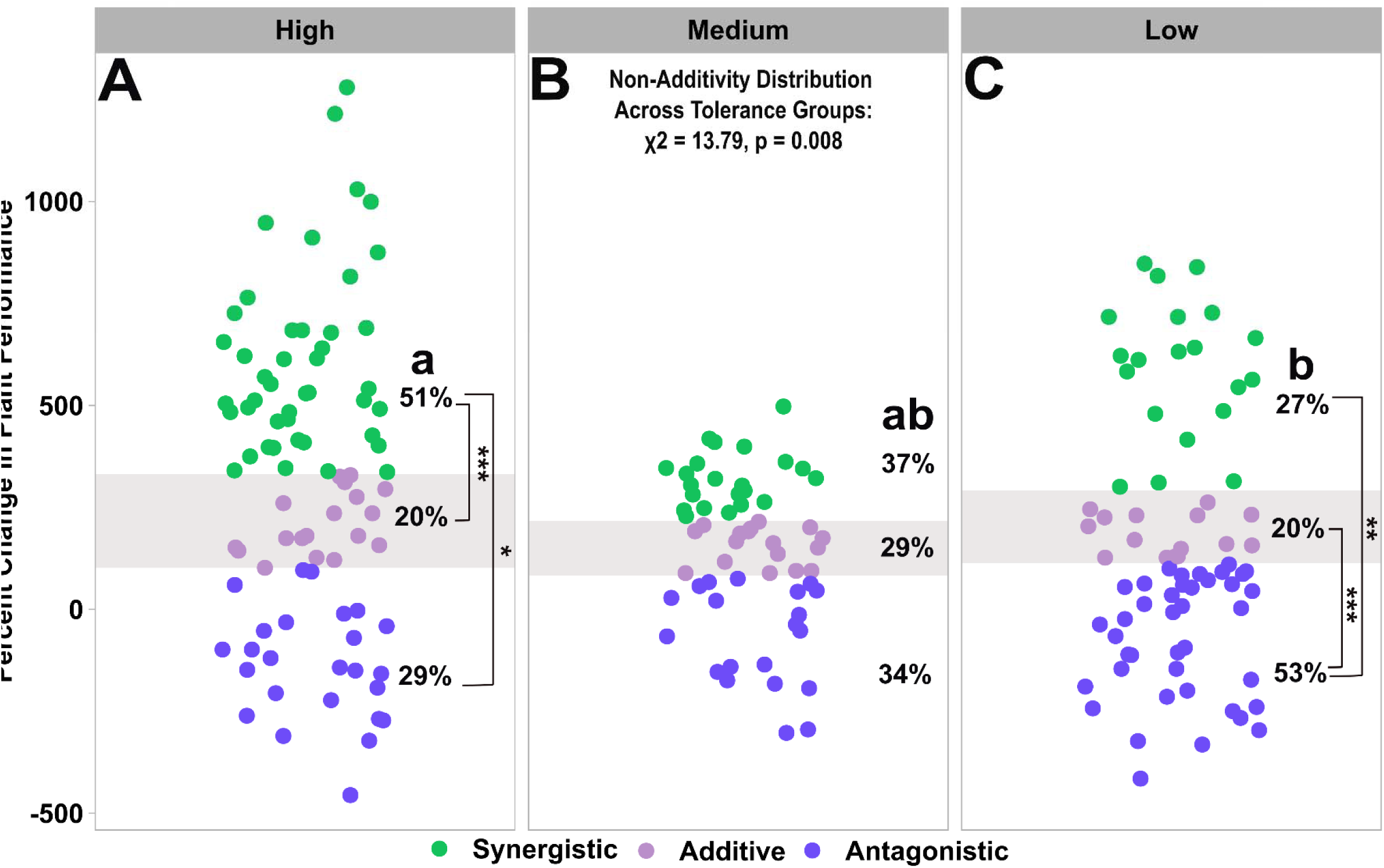
Non-additivity distributions of multiple mutualists on plant performance are significantly shaped by plant tolerance cluster where the majority of high tolerance plants display synergism while the majority of low tolerance plants display antagonism. Percent change in plant performance of plants for genotypes classified as having **(A)** high, **(B)** medium, or **(C)** low tolerance to general fungal pathogen invasion. Each point represents the change in plant performance of a biological replicate inoculated with multiple mutualists (R+M+) relative to controls with no mutualists (R-M-) measured in a benign environment at 5 weeks post-inoculation with mutualists. Percent change in plant performance axis values are shared across all panels. The gray shaded area is the 95% CI of the additive expectation. Points that fall within the gray area show additive effects (in purple), points that are above the gray area show synergistic effects (in green), and points that are below the gray area show antagonistic effects (in blue). Percentages represent the frequency of synergistic, additive, and antagonistic effects respectively. Statistics are provided for an overall X^2^ test of homogeneity to compare the non-additivity distributions among the three tolerance clusters, with different letters denoting tolerance cluster with significantly different distributions of non-additive outcomes determined by pairwise X^2^ post-hoc tests. Asterisks represent which non-additivity categorie are significantly different within a tolerance cluster where * is p<0.05, ** is p<0.01, and *** is p<0.001.

### Plant non-additivity and growth responses to multiple mutualists remain the same in a pathogenic environment for high and medium tolerance plants but decrease in low tolerance plants

For the high and medium tolerance clusters, plant non-additivity responses remained the same in both benign and pathogenic environments at 7 weeks post-inoculation with mutualists (chi-square Monte Carlo simulation; High: X^2^ = 4.72, p = 0.09, Fig. 3A; Medium: X^2^ = 0.40, p = 0.83, Fig. 3B) and was the same response as in the earlier benign timepoint of 5 weeks post-inoculation with mutualists (chi-square test of homogeneity; High: X^2^ = 1.49, p = 0.47, Fig. 2A; Medium: X^2^ = 4.01, p = 0.13, Fig. 2B). When comparing the change in plant growth through time; the high tolerance cluster grew on average 52% larger two weeks later in both the benign and pathogenic environments (repeated measures ANOVA: F_1,85_ = 16.21, p < 0.001; Fig. 4A; Table S2) demonstrating these plants were able to maintain the same rate of growth increase even when exposed to a pathogen. The size of the plants in the medium tolerance cluster remained constant through time in both environments (repeated measures ANOVA: F_1,57_ = 0.89, p = 0.35, Fig. 4B; Table S2). This indicates that for high and medium tolerance clusters, multiple mutualists confer the same non-additive performance and growth response outcomes in both benign and pathogenic environments suggesting that the presence of a pathogen does not shift the net benefits/costs the host experiences when associating with multiple mutualists. In contrast, the low tolerance cluster shifted to having significantly fewer synergistic responses in the pathogenic environment compared with the benign environment (chi-square Monte Carlo simulation: X^2^ = 7.15, p = 0.03; Fig. 3C). The low tolerance cluster grew 63% larger on average through time in benign conditions (repeated measures ANOVA: F_1,38_ = 8.19, p = 0.007; Fig. 4C; Table S2), but showed no growth through time when exposed to the pathogenic environment (repeated measures ANOVA: F_1,35_ = 0.50, p = 0.49; Fig. 4C; Table S2) indicating that exposure to a pathogen severely impacts their growth. These results suggest that the negative effect of multiple mutualists on plant performance within the low tolerance cluster is further exacerbated in a pathogenic environment.

**Fig. 3.**
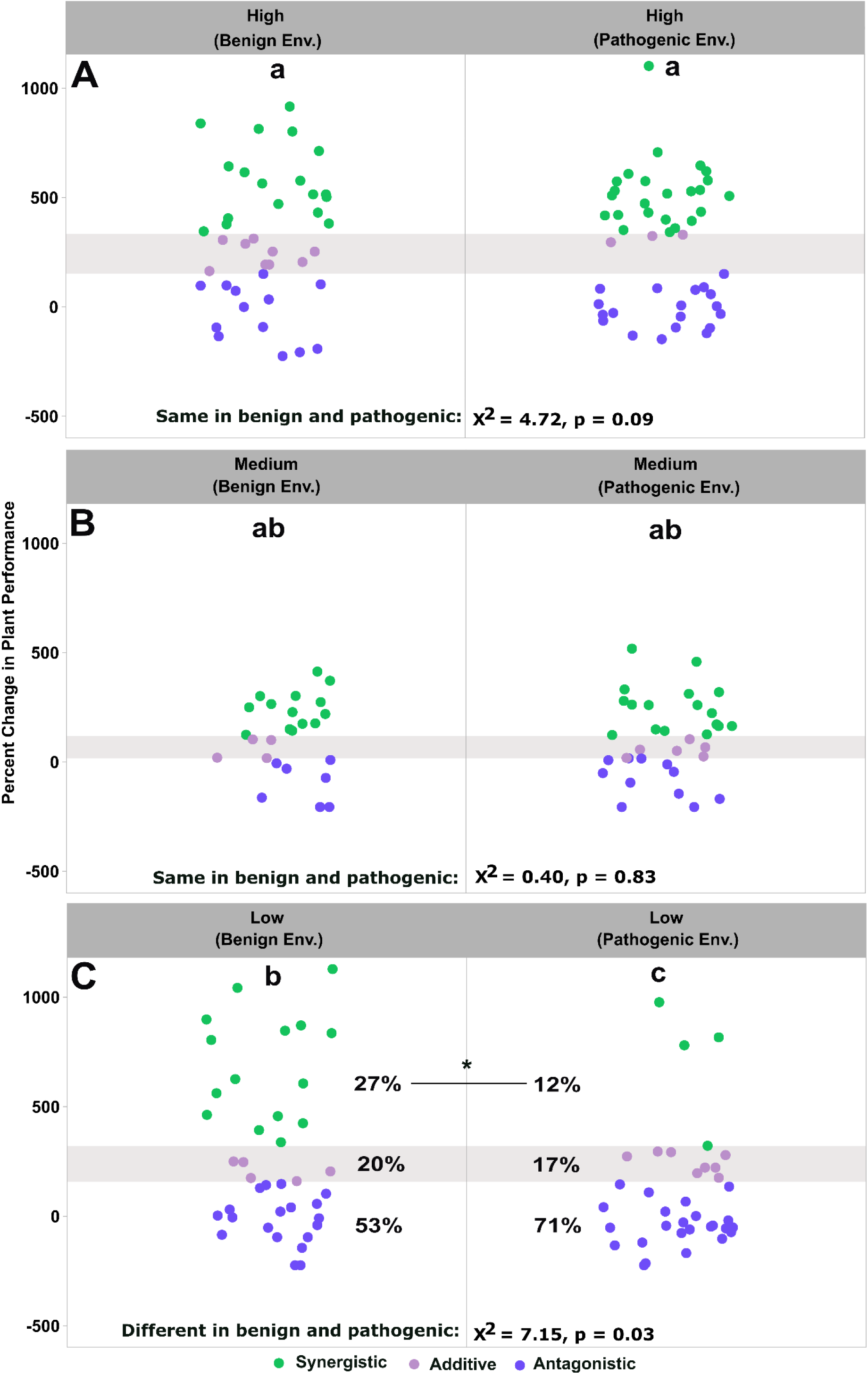
Plant non-additive responses remain the same in a pathogenic environment for high and medium tolerance clusters but have fewer synergistic responses in the low tolerance cluster. Percent change in plant performance of plants classified as having **(A)** high, **(B)** medium, or **(C)** low fungal pathogen tolerance. Plot pairs (left/right plots) contrast change in plant performance in a benign (left plot) and a pathogenic (right plot) environment. Each point represents the change in plant performance of a biological replicate inoculated with multiple mutualists relative to controls with no mutualists measured at 7 weeks post-inoculation with mutualists (i.e. 2 weeks post-inoculation with pathogen). Percent change in plant performance values are shared across all panels. The gray shaded area is the 95% CI of the additive expectation. Points that fall within the gray area show additive effects (in purple), points that are above the gray area show synergistic effects (in green), and points that are below the gray area show antagonistic effects (in blue). Stats are given for X^2^ Monte Carlo simulation tests between benign and pathogenic environments. Different letters indicate which tolerance clusters have significantly different non-additivity frequency distributions determined by pairwise X^2^ post-hoc tests among the tolerance clusters. The asterisk in panel C represents that the synergistic category within the low tolerance cluster is significantly different in a pathogenic environment where * is p<0.05 and the percentages represent the frequency distribution of synergistic, additive, and antagonistic effects respectively.

**Fig. 4.**
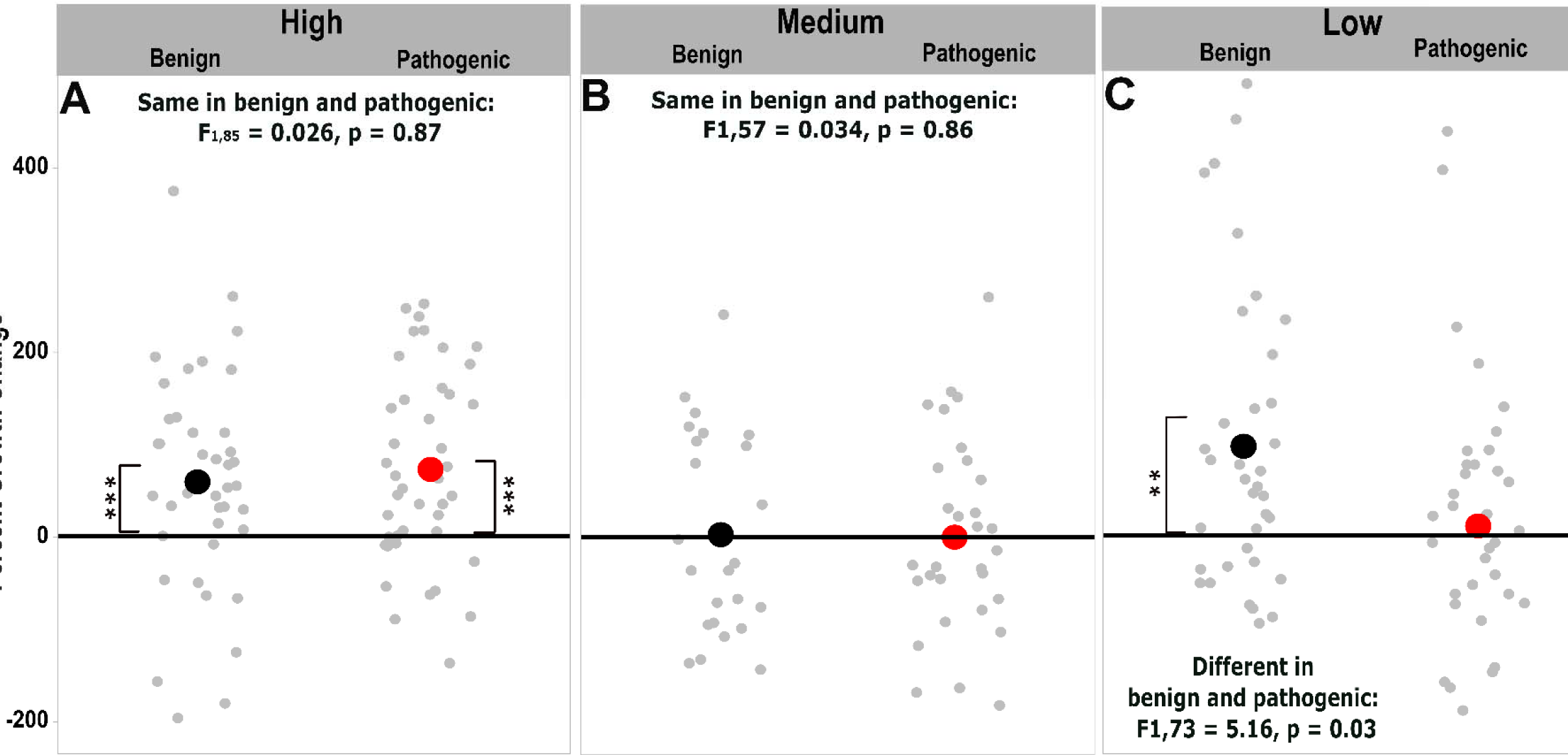
Plant growth responses remain the same in a pathogenic environment for high and medium tolerance plants but decrease in low tolerance plants. Percent change in plant growth of plants inoculated with multiple mutualists (R+M+) for genotypes classified as having **(A)** high, **(B)** medium, or **(C)** low tolerance to general fungal pathogen invasion. Percent growth change axis values are shared across all panels. Each gray point represents the change in plant growth of the same biological replicate measured at 5 weeks post-inoculation with mutualists and 7 weeks post inoculation with mutualists (i.e. 2 weeks post-inoculation with pathogens). Larger circles are the means of plant growth change for benign (black) and pathogenic (red) environments. Statistics are given for a repeated measures ANOVA comparing the means between benign and pathogenic environments. In addition, statistics are conveyed for a repeated measures ANOVA comparing the size of the plant across time. If the mean does not intersect the zero line, then the growth rate was found to be significantly different from zero denoted by ** p<0.01 and *** p<0.001.

### Low tolerance plants have significantly less AMF and symbiosis gene expression in multiple mutualist groups compared with single mutualist inoculation

The low tolerance cluster repeatedly showed decreased levels of AMF abundance and gene expression in the multiple mutualist group compared with the single mutualist groups in both pathogenic and benign environments. The low tolerance cluster had an 80% reduction in the abundance of AMF (Wilcoxon test: W = 756, p = 0.002, Fig. 5A) and around half the expression of PT8 phosphorous transport genes (Welch’s t-test: t_188.61_ = 1.96, p = 0.05, Fig 5B) in the treatment with both mutualists compared with the AMF only treatment. In addition, the low tolerance cluster had 60% less aquaporin (TC100851) expression, a gene associated with water/nutrient transport and disease regulation, in the treatment with both mutualists compared with the additive expectation based on single mutualist expression (ANOVA: F_2,106_ = 5.74, p = 0.004, Fig. 5C). Collectively, the lower AMF abundance, phosphorus transport expression, and aquaporin symbiosis quality in plants with multiple mutualists may contribute to this tolerance type mainly showing an antagonistic response to multiple mutualists.

**Fig. 5.**
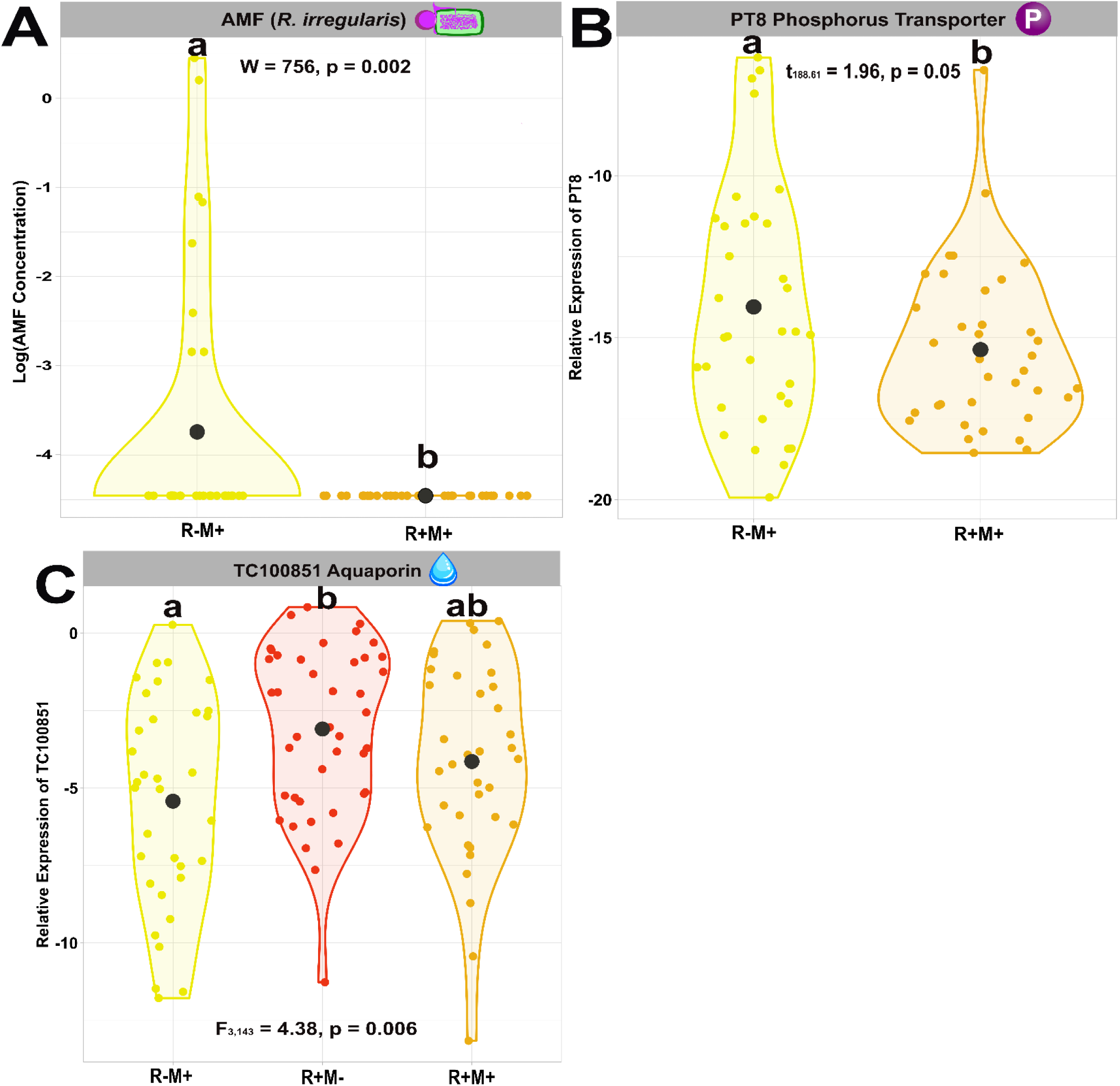
Low tolerance plants have significantly less AMF and symbiosis gene expression in multiple mutualist groups compared with single mutualist inoculation. Violin plots of **(A)** log AMF abundance, **(B)** relative expression of PT8 phosphorous transport gene, and **(C)** relative expression of TC100851 aquaporin gene. Each small point represents data obtained from a biological replicate while the larger black circles demarcate the mean. Each color indicates the mutualist inoculation status; yellow are plants with AMF being the only mutualist, red are plants with rhizobia being the only mutualist, and orange are plants with both AMF and rhizobia. Abundance data is on a log10 scale while relative gene expression data is on a log2 scale. Wilcoxon, Welch t, and ANOVA statistics are provided respectively to show significant differences between groups. Different letters signify groups that ar significantly different from one another using Tukey post-hoc tests.

### Plants with synergistic growth responses have more rhizobia associations and higher quality mutualisms than plants with antagonistic growth responses

Within the multiple mutualist group, rhizobia abundance and symbiosis gene expression levels corresponded to the non-additivity category of the host plant. Host plants that displayed synergistic growth responses to multiple mutualists had over 600% higher rhizobia abundance than plants with antagonistic responses while plants with additive responses showed intermediate rhizobia abundance (ANOVA: F_2,107_ = 3.96, p = 0.02; Fig 6A). Both plants with synergistic and additive responses had significantly higher expression of genes indicative of excellent symbiosis quality, with ∼140% higher aquaporin TC100851 (ANOVA: F_2,104_ = 3.48, p = 0.03; Fig 6B), 200% higher phosphorus transporter PT8 (ANOVA: F_2,104_ = 3.76, p = 0.03; Fig 6C), and 200% higher ammonium transporter AMT2;3 expression (ANOVA: F_2,104_ = 6.20, p = 0.003; Fig 6D) than host plants that displayed antagonistic responses regardless of whether the environment was benign or pathogenic. These multiple lines of evidence indicate that plants with synergistic growth responses have more rhizobia associations and higher quality mutualisms than that of plants with antagonistic growth responses. Since there are disproportionately higher rates of synergism and antagonism in high and low tolerance genotypes respectively, these results suggest that plant genotype plays a critical role in modulating microbial associations and mutualism quality within a species and implies that some of the same mechanisms that allow plants to regulate pathogens are important in the regulation of symbionts. Interestingly, fusarium concentration was the same across all tolerance clusters, treatment groups, and non-additivity categories tested (ANOVA Group x Tolerance: F_6,190_ = 0.27, p = 0.95; ANOVA Non-additivity Category: F_2,55_ = 0.23, p = 0.80; Fig. S3). This result suggests that high tolerance clusters may not be functionally impacted as severely by a high pathogen load or may be more effective at clearing the pathogen after infection has established, rather than preventing the pathogen from replicating to a high load.

**Fig. 6.**
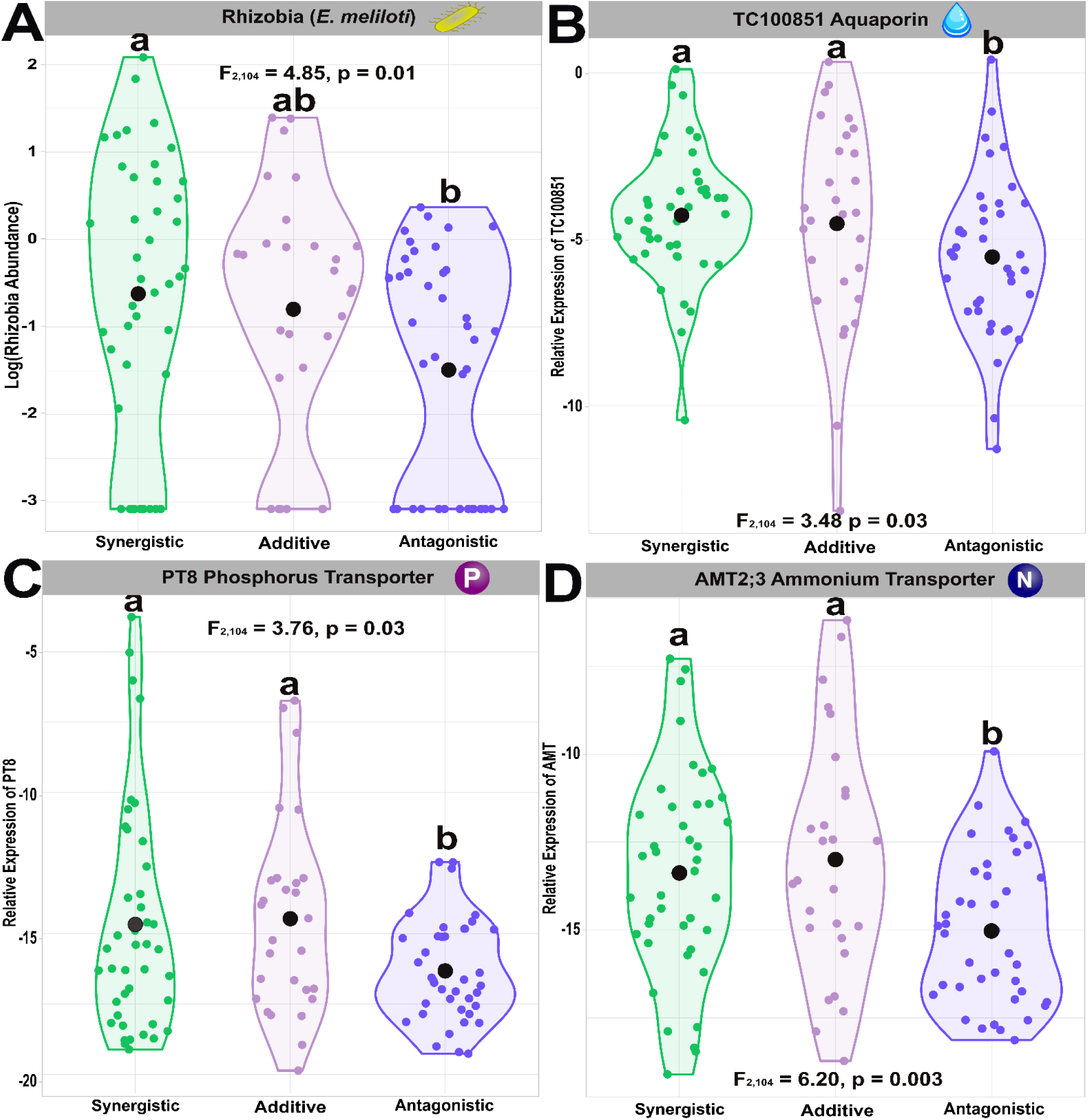
Plants with synergistic growth responses have more rhizobia associations and higher quality mutualisms than plants with antagonistic growth responses. Violin plots of **(A)** log rhizobia abundance, **(B)** relative expression of TC100851 aquaporin gene, **(C)** relative expression of PT8 phosphorous transport gene, and **(D)** relative expression of AMT2:3 ammonium transporter gene. Each small point represents data obtained from a biological replicate while the larger black circles demarcate the mean. Each color indicates the plant non-additivity response to multiple mutualists; green are plants with a synergistic response, purple are plants with an additive response, and blue are plants with an antagonistic response. Abundance data is on a log10 scale while relative gene expression data is on a log2 scale. ANOVA statistics are provided to show significant differences between groups. Different letters signify groups that are significantly different from one another using Tukey post-hoc tests.

### Plant survival against pathogen is dependent upon host tolerance genotype and mutualist-mediated plant performance

Plants with a higher composite plant performance score (i.e. larger, more robust plants) prior to pathogen invasion were most likely to survive pathogen stress (Logistic Regression: z = 5.17, p < 0.001; Fig. 7A). In pathogenic environments, high and medium tolerance clusters inoculated with multiple mutualists were the largest in size (Fig. 7B,D) and showed the highest survival rates relative to controls with no mutualists (∼35% higher survival; Fig. 7C,E). Plants with rhizobia as the only mutualist were intermediate/smaller in size (Fig. 7B,D) and had an intermediate survival increase of ∼20% higher survival (Fig. 7C,E). Plants with AMF as the only mutualist were similar in size to controls (Fig. 7B,D) and showed similar survival rates to controls (Fig. 7C,E) indicating that AMF alone was not enough to increase plant survival, potentially due to nitrogen being a limiting factor. In contrast, in the low tolerance cluster, plants inoculated with multiple mutualists or just AMF were similar in size to controls (Fig. 7F) and only had a slight increase in survival relative to controls (∼10% higher survival; Fig. 7G). However, the low tolerance plants were generally larger (Fig. 7F) and had significantly higher survival when only inoculated with rhizobia (∼50% higher survival; Fig. 7G). Overall, these results indicate that survival in the pathogenic environment is largely dependent upon plant performance prior to pathogen invasion, since larger and healthier plants had the best survival rates. Therefore, the interaction between host tolerance genotype and mutualist status not only significantly impacted plant performance, but also significantly impacts plant fitness when exposed to fungal pathogen stress. These results further corroborate that hosts with multiple mutualists perform better than hosts with single mutualists only in high and medium tolerance genotypes, while in low tolerance genotypes hosts with multiple mutualists perform worse than hosts with only rhizobia. This again shows that high and medium tolerance genotypes experience a net benefit in fitness when associating with multiple mutualists while low tolerance genotypes experience a net cost.

**Fig. 7.**
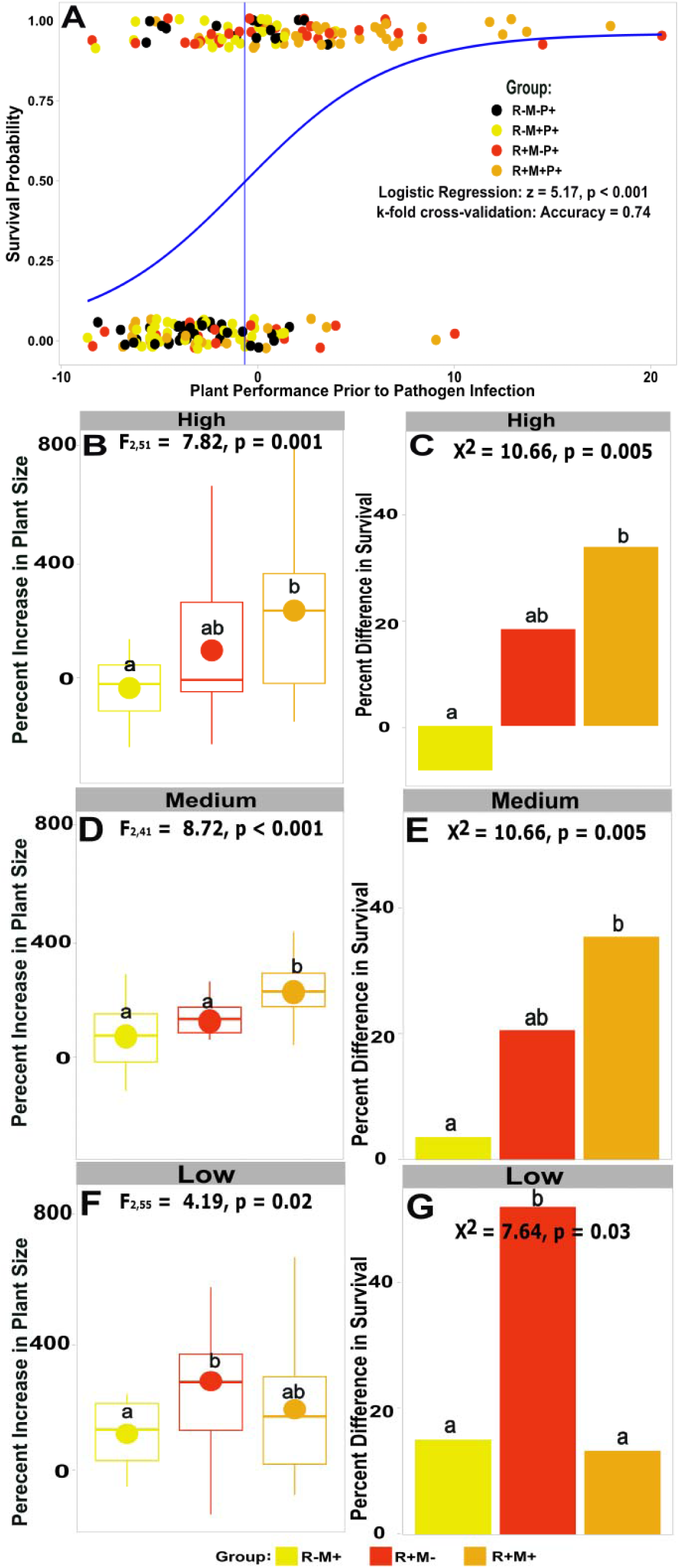
Plant survival against pathogen is dependent upon host tolerance type and mutualist-mediated plant performance. **(A)** Logistic regression of pathogen survival predicted by composite plant performance metric prior to pathogen inoculation. Statistics are provided for the logistic regression and k-fold cross-validation accuracy of the model (Survival ∼ Plant Performance). Percent increase in plant size prior to pathogen inoculation and percent difference in pathogen survival relative to controls with no mutualists of plants for genotypes classified as having **(B)(C)** high, **(D)(E)** medium, or **(F)(G)** low tolerance to general fungal pathogen invasion. Statistics indicate significant differences in plant size or survival using ANOVAs and X^2^ proportion tests respectively. Different letters signify groups that are significantly different from one another using Tukey post-hoc and Fisher pairwise tests respectively. The color of the points/bars denotes the mutualist inoculation status; yellow are plants with AMF being the only mutualist, red are plants with rhizobia being the only mutualist, orange are plants with both AMF and rhizobia, and black are controls with no mutualists.

## Discussion

Rhizobia and AMF symbioses are considered to be among the most important symbioses on Earth due to their diverse roles in ecosystem and agricultural health and resilience (Tian, Khah, and Kariman 2019). Rhizobia and AMF are also the major microbial supplements used for legume agricultural soil improvement where they provide substantial nutritional benefits while being a cheaper, eco-friendly alternative to chemical fertilizers (George and Ray 2023, Kebede 2021, Kumar, Sindhu, and Kumar 2022). However, microbial supplements have faced challenges with low efficacy as they can be difficult to reliably establish in field conditions (Kaminsky et al. 2019) and often do not have the same effects on plant performance in the field as in the lab due to the combination of nonadditive interactions with other microbes, modulation of symbioses effects in a pathogenic environment, and diversity of differing plant genotypic responses to inoculation (Maged, Abdul, and Heribert 2020). In order to successfully harness the benefits of microbial symbioses, we need a comprehensive understanding of the complex interplay between plant genotype, multiple mutualists, and benign/pathogenic environmental conditions on plant performance. Our study fills this knowledge gap by combining a large-scale manipulative experiment, plant performance/survival analyses, RNA extractions, and targeted gene expression to comprehensively assess how multiple mutualists interact with host tolerance genotype to shape the severity of pathogenic disease. Our results reveal three important findings: (1) host genotypes with high tolerance to pathogens benefit more from multiple mutualist interactions than susceptible genotypes, (2) microbial mutualists confer the same plant performance benefits in both benign and pathogenic environments, and (3) the quality of the symbiotic relationship with multiple mutualists is a strong predictor of survival against fungal pathogenic attack. Below we discuss how the results from this study both advance our understanding of plant disease ecology and inform new directions for identifying what genetic features of a legume may simultaneously enable high quality symbioses while conferring tolerance to fungal pathogens.

Symbioses between plants and “microbes-of-large-effects” can shift from mutualistic to parasitic depending on the abiotic and biotic environmental contexts (Afkhami et al. 2020). One important biotic factor is the presence of other microbes and their non-additive interactions where microbes facilitating or competing with one another can lead to synergistic and antagonistic effects respectively (Afkhami et al. 2020, Albornoz et al. 2022). For example, one study showed when co-inoculating flax with fungal isolates differing in their carbon sink strength, the fungi facilitated one another’s growth and provided synergistic complementary rewards to the host plants enabling them to both grow rapidly and maintain a large plant biomass (Martignoni et al. 2021). Another study showed that competition between fungal endophytes and mycorrhizae in grasses reduced one another’s abundances and negatively impacted host plant growth and blade yield (Liu et al. 2011). A recent paper found that interactions within more than half of experimental fungal consortia synergistically or antagonistically affected morning glory host plant performance, sometimes with extreme consequences (e.g. >1000% increase in host productivity above the additive expectation; Almeida, Tran, and Afkhami 2024). Similar to these studies, our results indicated that when the plant performance outcome was synergistic, microbes were facilitating one another’s growth (600% increase in rhizobia) and likely providing complementary rewards to the host (upregulated inducement of host nutrient receptors involved in resource acquisition from symbionts: PT8 phosphorus, AMT2;3 ammonium, and TC10085 aquaporin) compared with antagonistic interactions. In addition to antagonistic interactions performing worse in these areas, we also found further evidence these microbes were competing and reducing one another’s abundances (80% less AMF) and reduced rewards (∼60% less aquaporin) by comparing against their single mutualist outcomes.

Host genotype has also been shown to be an influential biotic factor in determining microbial recruitment and different genotypes of plants within the same species often differ in their ability to recruit microbes with high quality benefits (Hohmann and Messmer 2017). In recent years, it has been demonstrated across many diverse plant species including sugar cane (Ishida et al. 2022), Monterey pine (Gallart et al. 2018), soybean (Zhong et al. 2019), tomato (Cordovez et al. 2021), Balsam poplar (Rheault et al. 2020), and sunflowers (Bueno de Mesquita et al. 2024) that host genotype is a critical component of rhizosphere assembly. However, it is less clear the extent that host genotype modulates intermicrobial interactions once microbes are recruited.

Importantly, our study demonstrated that the likelihood of a legume receiving synergistic, additive, or antagonistic plant growth outcomes was largely dependent upon host genotype modulation of multiple mutualists. Since the strains of rhizobia and AMF and abiotic environmental factors were kept consistent, we therefore know that genotype must have been a key factor in the microbial interactions switching from facilitation to competition.

Excitingly, we found that genotypes that have high tolerance to fungal root pathogens also had mostly synergistic responses to multiple mutualists while the low tolerance genotypes had mostly antagonistic responses. Intriguingly, our findings suggest that genotypes that fall within the same pathogen tolerance cluster have shared genetic characteristics that make them not only similar in their tolerance to fungal pathogens but also similar in how they modulate multiple mutualist interactions. One possible hypothesis documented in plants is that hosts that get synergistic benefits from mutualists have better nutrition, growth, and overall health making them more robust against pathogen infection (Dordas 2008, Rahman et al. 2018). Another possibility is that the host plant, by promoting microbial facilitation, helps to maximize mutualist abundance.

Higher mutualist abundances often reduce a pathogen’s access to the host (Sassone-Corsi 2015) since the mutualists are physically occupying a higher proportion of the available root surface area. In terms of biochemical pathways, there could also potentially be differences among genotypes in how they regulate the common symbiosis pathway such as differential activation of calcium- and calmodulin-dependent kinase (CCaMK) which in addition to modulating rhizobial and AMF symbioses in plants has also been shown to play a role in plant responses to pathogenic fungi (Genre and Russo 2016).

Surprisingly, we also discovered that in high tolerance hosts, microbial mutualists provided the same high nutritional (high expression levels of host nutrient receptors), plant performance (∼500% better than controls), and growth rate (50% size increase through time) benefits to plants in both the benign and pathogenic environments. This is noteworthy because there are often significant trade-offs between plant growth and pathogen defense strategies (He, Webster, and He 2022, Huot et al. 2014), where manipulations of plant genes favoring growth result in a suppressed immune response and vice versa. Recently, it has been suggested that microbiomes may be the key to mitigating this growth-defense trade-off since they can simultaneously promote plant growth and decrease plant pathogen susceptibility (Karasov et al. 2017, Ku et al. 2024). Our study further supports this hypothesis since our survival analyses revealed that legumes that facilitated high quality and synergistic mutualism interactions both grew larger and had higher survival. This finding has important consequences for agricultural breeding since both nutrition and pathogen defense are crucial for crop success. By developing legume crops that impose stronger selection on microbes for greater mutualism potential, crop yields and pathogen defense could be simultaneously increased while reducing the need for toxic fertilizers and pesticides (Denison and Muller 2022).

We also found that low tolerance hosts had the same low nutritional status in both environments (low expression levels of host nutrient receptors), and even worse plant performance (∼100% worse than controls) and growth rate (reduced 60% through time) in the pathogenic environment. Since they were reduced in size and the size of the plant was a significant predictor of survival, most of the low tolerance hosts also died before reaching maturation for production of seed pods. Even though these low tolerance hosts performed the worst in terms of interactions with symbionts, symbiont-mediated growth benefits, and survival, these genotypes still exist in nature (Lei et al. 2019), indicating that they must still exhibit some type of fitness advantage under at least some conditions. We postulate that these genotypes are likely to produce a higher quantity of seed pods to compensate for their reduced growth and survival. However, we harvested our plants for the final data collection at the onset of first seed production (where plants were considered to have successfully survived the pathogen) so were not able to collect total pods produced. There is some evidence for a trade-off between high quality associations with mutualists and seed production (Gundel et al. 2012) and seed viability (Fuchs et al. 2024) in cool-season grass species, but this topic has remained largely understudied. Therefore, further work is needed to assess whether seed production is attenuated in legumes that have high quality associations with multiple mutualists and if this is a common trade-off among diverse plant species.

Our study also opens up other new areas for identifying what genetic features of a legume may simultaneously enable high quality symbioses while conferring tolerance to fungal pathogens. We found that plant genotype was a significant factor in modulating whether multiple mutualists acted synergistically or antagonistically, with genotypes highly tolerant to pathogens best able to maximize benefits and minimize costs of interacting with multiple mutualists. However, the genes responsible for these differences remain unknown. We suggest that future work could use genome-wide association studies and knock down of candidate genes in tandem to identify host plant genes that simultaneously underpin regulation both of complex mutualist interactions and pathogen effects. We also found that survival against pathogen attack was predicted by the robustness in growth of the legume prior to pathogen invasion. By using whole transcriptome sequencing technologies such as 3’RNA-seq, we could determine whether additional plant immunity pathways such as hormonal signaling of jasmonic-ethylene/salicylic acid (Li et al. 2019) and saponin/volatile oil production (Lei et al. 2019) are also modified through differential interactions with mutualists in a pathogenic environment. Overall, our study lays the groundwork for better understanding the complex interaction among host genotype, microbial mutualists, and pathogens necessary for legume agricultural improvement.

## Data Availability

All raw data, supporting data, and scripts used to produce this manuscript is available open access in Zenodo (https://doi.org/10.5281/zenodo.14569960).

## Acknowledgements

We thank J. Prosperi (INRA-Montpellier) and L. Gentzbittel (ENSAT-Toulouse) for providing the original legume source material via the Medicago HapMap project and N. Young (University of Minnesota) for consultation on the germplasm and genetic resources available for the Medicago HapMap project. We appreciate the National Center for Genome Resources (Santa Fe, NM, USA) and the USDA-ARS Legume Germplasm Repository (Pullman, WA, USA) for supplying additional *M. truncatula* seeds. We also thank D. Samac at the USDA-ARS Plant Science Research Unit (St. Paul, MN, USA) for supplying the *Fusarium oxysporum f.sp. medicaginis* under USDA-APHIS permit P526-21-05187. We are also grateful to K. Mayol-Graciano, Y. Quevedo, and Y. Chen. for assistance in plant growth and maintenance and the C. Searcy lab for feedback on the manuscript. Research was funded by University of Miami research funds awarded to M. Afkhami and the Mycological Society of America graduate fellowship awarded to A. Rawstern. A. Rawstern was additionally supported by the University of Miami Department of Biology and the Lisa D. Anness Graduate Fellowship.

## References

Afkhami, M., Rudgers, J., and Stachowicz, J. Multiple mutualist effects: conflict and synergy in multispecies mutualisms. Ecology. 95, 833–844 (2014).

Afkhami, M. and Stinchcombe, J. (2016). Multiple mutualist effects on genomewide expression in the tripartite association between *Medicago truncatula*, nitrogen-fixing bacteria and mycorrhizal fungi. Mol. Ecol. 25, 4946–4962 (2016).

Afkhami, M., Almeida, B., Hernandez, D., Kiesewetter, K., Revillini, D. Tripartite mutualisms as models for understanding plant–microbial interactions. Curr. Opin. Plant Biol. 56, 28– 36 (2020).

Afkhami, M., Friesen, M., and Stinchcombe, J. Multiple mutualism effects generate synergistic selection and strengthen fitness alignment in the interaction between legumes, rhizobia, and mycorrhizal fungi. Ecol. Lett. 24, 1824–1834 (2021).

Albornoz, F., Prober, S., Ryan M., Standish R. Ecological interactions among microbial functional guilds in the plant-soil system and implications for ecosystem function. Plant Soil. 476, 301–313 (2022).

Almeida, B., Tran, E., and Afkhami, M. Phyllosphere fungal diversity generates pervasive nonadditive effects on plant performance. New Phytol. 19792 (2024).

Anderson, P., Cunningham, A., Patel, N., Morales, F., Epstein, P., Daszak, P. Emerging infectious diseases of plants: pathogen pollution, climate change and agrotechnology drivers. TREE. 19, 535–544 (2004).

Ankita, Choudhary, S., Bakala, H., Sarao, L., Kaur, S. (2023). Pulses Waste to Biofuels. In: Srivastava, N., Verma, B., Mishra, P. (eds) Agroindustrial Waste for Green Fuel Application. Clean Energy Production Technologies. Springer, Singapore.

Baichman-Kass, A., Song T., and Friedman, J. Competitive interactions between culturable bacteria are highly non-additive. eLife. 12, e83398 (2023).

Barker, D. G., Pfaff, T., Moreau, D., Groves, E., Ruffel, S., Lepetit, M., Whitehand, S., Maillet, F., Nair, R. M., and Journet, E.-P. (2006). Growing M. truncatula: Choice of substrates and growth conditions. In: The Medicago truncatula Handbook (eds Mathesius U., Journet EP, Sumner LW) Noble Foundation, Ardmore.

Benedito, V., Li, H., Dai, X., Wandrey M., He, J., Kaundal, R., et al. Genomic inventory and transcriptional analysis of *Medicago truncatula* transporters. Plant Physiol. 152, 1716–1730 (2010).

Bisht, A., Sharma, V., Garg, N. (2024). Deciphering the Role of Arbuscular Mycorrhizal Fungi in Mitigating the Negative Effects of Abiotic Stresses in Legume Crops. In: Parihar, M., Rakshit, A., Adholeya, A., Chen, Y. (eds) Arbuscular Mycorrhizal Fungi in Sustainable Agriculture: Nutrient and Crop Management. Springer, Singapore.

Boyd, I., Freer-Smith P., Gilligan, C., Godfray H. The consequence of tree pests and diseases for ecosystem services. Science. 342, 1235773 (2013).

Breuillin-Sessoms, F., Floss, D., Gomez, S., Pumplin, N., Ding, Y., Levesque-Tremblay, V. et al. Suppression of arbuscule degeneration in *Medicago truncatula* phosphate transporter4 mutants is dependent on the ammonium transporter 2 family protein AMT2;3. The Plant Cell. 27, 1352–1366 (2015).

Bueno de Mesquita, C., Walsh, C., Attia, Z., Koehler, B., Tarble, Z., Van Tassel, D., et al. Environment, plant genetics, and their interaction shape important aspects of sunflower rhizosphere microbial communities. Appl. Environ. Microbiol. 90, 01635 (2024).

Bulgarelli D., Schlaeppi K., Spaepen, S., Ver Loren van Themaat, E., Schulze-Lefert. Structure and functions of the bacterial microbiota of plants. Annu. Rev. Plant Biol. 64, 807–38 (2013).

Chabaud, M., Harrison, M., de Carvalho-Niebel, F., Bécard, G., and Barker, D. G. (2006). Inoculation and Growth with Mycorrhizal Fungi. In: The Medicago truncatula Handbook (eds Mathesius U., Journet EP, Sumner LW) Noble Foundation, Ardmore.

Chen, X., Koumoutsi, A., Scholz, R., Schneider, K., Vater, J, Süssmuth, R. et al. Genome analysis of *Bacillus amyloliquefaciens* FZB42 reveals its potential for biocontrol of plant pathogens. J. Biotechnol. 140, 27–37 (2009).

Choudhury, R., Er, H., Hughes, M., Smith, J., Pruett, G., Konkol, J. et al. Host density dependence and environmental factors affecting laurel wilt disease incidence. Plant Pathol. 70, 676–688 (2021).

Chuanen, Z., Han, L., Pislariu, C., Nakashima, J., Fu, C., Jiang, Q. et al. From model to crop: functional analysis of a STAY-GREEN gene in the model legume *Medicago truncatula* and effective use of the gene for alfalfa improvement. Plant Physiol. 157, 1483–1496 (2011).

Cope, K., Kafle, A., Yakha, J., Pfeffer, P., Strahan, G., Garcia, K., et al. Physiological and transcriptomic response of *Medicago truncatula* to colonization by high- or low-benefit arbuscular mycorrhizal fungi. Mycorrhiza. 32, 281–303 (2022).

Cordovez, V., Rotoni, C., Dini-Andreote, F., Oyserman, B., Carrión, V., Raaijmakers, J. Successive plant growth amplifies genotype-specific assembly of the tomato rhizosphere microbiome. Sci. Total. Environ. 772, 144825 (2021).

Cycoń, M., Mrozik, A., and Piotrowska-Seget, Z. Antibiotics in the soil environment— degradation and their impact on microbial activity and diversity. Front. Microbiol. 10, 338 (2019).

Denison, R., and Muller, K. An evolutionary perspective on increasing net benefits to crops from symbiotic microbes. Evol. Appl. 15, 1490–1504 (2022).

De Bang, T., Lundquist, P., Dai, X., Boschiero, C., Zhuang, Z., Pant. P., et al. Genome-wide identification of Medicago peptides involved in macronutrient responses and nodulation. Plant Physiol. 175, 1669–1689 (2017).

De Souza, R., Seibert, D., Quesada, H., Bassetti, F., Fagundes-Klen, M., Bergamasco, R. Occurrence, impacts and general aspects of pesticides in surface water: a review. PSEP. 135, 22–37 (2020).

Dong, O., and Ronald, P. Genetic engineering for disease resistance in plants: recent progress and future perspectives. Plant Physiol. 180, 26–38 (2019).

Dordas, C. Role of nutrients in controlling plant diseases in sustainable agriculture. A review. Agron. Sustain. Dev. 28, 33–46 (2008).

Esse, H., Reuber, T., and Does, D. Genetic modification to improve disease resistance in crops. New Phytol. 225, 70–86 (2020).

Fleige, S., Walf, V., Huch, S., Prgomet, C., Sehm, J., Pfaffl, M. Comparison of relative mRNA quantification models and the impact of RNA integrity in quantitative real-time RT-PCR. Biotechnol Lett. 28, 1601–1613 (2006).

Fuchs B., Damerau, A., Yang, B., Muola, A. Reduced seed viability in exchange for transgenerational plant protection in an endophyte-symbiotic grass: does the defensive mutualism concept pass the fitness test? Ann. Bot. mcae133 (2024).

Gallart, M., Adair, K., Love, J., Meason, D., Clinton, P., Xue, J. et al. Host genotype and nitrogen form shape the root microbiome of *Pinus radiata*. Microb. Ecol. 75, 419–433 (2018).

Garcia, J., Barker, D. G., and Journet, E.-P. (2006). Seed storage and germination. In: The Medicago truncatula Handbook (eds Mathesius U., Journet EP, Sumner LW) Noble Foundation, Ardmore.

Genre, A. and Russo, G. Does a Common Pathway Transduce Symbiotic Signals in Plant– Microbe Interactions? Front. Plant Sci. 7, 96 (2016).

George, N. and Ray, J. The inevitability of arbuscular mycorrhiza for sustainability in organic agriculture—A critical review. Front. Sustain. Food Syst. 7, 1124688 (2023).

Gordon, T. *Fusarium oxysporum* and the fusarium wilt syndrome. Annu. Rev. Phytopathol. 55, 23–39 (2017).

Gou, J., Debnath, S., Sun, L., Flanagan, A., Tang, Y., Jiang, Q. et al. From model to crop: Functional characterization of SPL8 in *M. truncatula* led to genetic improvement of biomass yield and abiotic stress tolerance in alfalfa. Plant Biotechnol. J. 16, 951–962 (2018).

Gundel P., Garibaldi, L., Martínez-Ghersa, Ghersa, C. Trade-off between seed number and weight: Influence of a grass-endophyte symbiosis. Basic Appl. Ecol. 13, 32–39 (2012).

Franklin, J., Hockey, K., and Maherali, H. Population-level variation in host plant response to multiple microbial mutualists. Am. J. Bot. 107, 1389–1400 (2020).

He, Z., Webster S., He, S. Growth-defense trade-offs in plants. Curr. Biol. 32, 634–639 (2022).

Hodge, A. Microbial ecology of the arbuscular mycorrhiza. FEMS Microbiol. Ecol. 32, 91–96 (2000).

Hohmann, P. and Messmer, M. Breeding for mycorrhizal symbiosis: focus on disease resistance. Euphytica. 213, 113 (2017).

Huot, B., Yao, J., Montgomery, B., He, S. Growth-defense tradeoffs in plants: a balancing act to optimize fitness. Mol. Plant. 7, 1267–1287 (2014).

Ishida, J., Bini, A., Creste, S., Van Sluys, M. Towards defining the core Saccharum microbiome: input from five genotypes. BMC Microbiol. 22, 193 (2022).

Ivashuta, S., Liu, J., Liu, J., Lohar, D., Haridas, S., Bucciarelli, B., et al. RNA interference identifies a calcium-dependent protein kinase involved in Medicago truncatula root development. The Plant Cell. 17, 2911–2921 (2005).

Journet, E.-P., de Carvalho-Niebel, F., Andriankaja, A., Huguet, T., and Barker, D. G. (2006). Rhizobial inoculation and nodulation of Medicago truncatula. In: The Medicago truncatula Handbook (eds Mathesius U., Journet EP, Sumner LW) Noble Foundation, Ardmore.

Kaminsky, L., Trexler, R., Malik, R., Hockett, K., Bell, T. The inherent conflicts in developing soil microbial inoculants. Trends Biotechnol. 37, 140–151 (2019).

Karagiannidis, N., Bletsos, F., and Stavropoulos, N. Effect of Verticillium wilt (*Verticillium dahliae* Kleb.) and mycorrhiza (*Glomus mosseae*) on root colonization, growth and nutrient uptake in tomato and eggplant seedlings. Sci. Hortic. 94, 145–156 (2002).

Karasov, T., Chae, E. Herman, J., Bergelson, J. Mechanisms to mitigate the trade-off between growth and defense. The Plant Cell. 29, 666–680 (2017).

Kasanke, S., Cheeke, T., Moran, J., Roley, S. Tripartite interactions among free-living, N-fixing bacteria, arbuscular mycorrhizal fungi, and plants: mutualistic benefits and community response to co-inoculation. SSSA. saj2.20679 (2024).

Kebede, E. Competency of rhizobial inoculation in sustainable agricultural production and biocontrol of plant diseases. Front. Sustain. Food Syst. 5, 728014 (2021).

Kinkel, L., Schlatter, D., Bakker, M., Arenz, B. Streptomyces competition and co-evolution in relation to plant disease suppression. Res. Microbiol. 163, 490–499 (2012).

Krol, E. and Becker, A. Global transcriptional analysis of the phosphate starvation response in *Sinorhizobium meliloti* strains 1021 and 2011. Mol. Genet. Genom. 272, 1–17 (2004).

Ku, Y., Liao, Y., Chiou, S., Lam, H., Chan, C. From trade-off to synergy: microbial insights into enhancing plant growth and immunity. Plant Biotechnol. J. 22, 2461–2471 (2024).

Kumar, S., Rajendran, K., Kuman, J., Hamwieh, A., Baum, M. Current knowledge in lentil genomics and its application for crop improvement. Front. Plant Sci. 6, 00078 (2015).

Kumar, D., Sindhu, S., and Kumar, R. Biofertilizers: An ecofriendly technology for nutrient recycling and environmental sustainability. Curr. Res. Microb. Sci. 3, 100094 (2022).

Langsrud, O. ANOVA for unbalance designs: Use type II instead of type III sum of squares. Stat. Comput. 13, 163–167 (2003).

Lei, Z., Watson, B., Huhman, D., Yang, D., Sumner, L. Large-scale profiling of saponins in different ecotypes of *Medicago truncatula*. Front. Plant Sci. 10, 850 (2019).

Leveau, J. Re-envisioning the plant disease triangle by integration of host microbiota and a pivot in focus to health outcomes. Annu. Rev. Phytopathol. 62, 10.1146 (2024).

Libault, M., Joshi, T., Benedito, V., Xu, D., Udvardi, M., Stacey, G. Legume transcription factor genes: what makes legumes so special? Plant Physiol. 151, 991–1001 (2009).

Lichtenzveig, J., Anderson, J., Thomas, G., Oliver, R., and Singh, K. (2006). Inoculation and growth with soil borne pathogenic fungi. In: The Medicago truncatula Handbook (eds Mathesius U., Journet EP, Sumner LW) Noble Foundation, Ardmore.

Li, N., Han X., Feng D., Yuan D., Huang L. Signaling crosstalk between salicylic acid and ethylene/jasmonate in plant defense: do we understand what they are whispering? Int J Mol Sci. 20, 671 (2019).

Liu, J., Maldonado-Mendoza, I., Lopez-Meyer M., Cheung, F., Town, C., Harrison, M. Arbuscular mycorrhizal symbiosis is accompanied by local and systemic alterations in gene expression and an increase in disease resistance in the shoots. TPJ. 50, 529–544 (2007).

Liu, Q., Parsons, A., Xue, H., Fraser, K., Ryan, G., Newman J. et al. Competition between foliar *Neotyphodium lolii* endophytes and mycorrhizal Glomus spp. fungi in *Lolium perenne* depends on resource supply and host carbohydrate content. Funct. Ecol. 25, 910–920 (2011).

Livak K. and Schmittgen, T. Analysis of relative gene expression data using real-time quantitative PCR and the 2(-Delta Delta C(T)) method. Methods. 25, 402–408 (2001).

Lodwig, E., Hosie, A., Bourdès, A., Findlay, K., Allaway, D., Karunakaran, R. et al. Amino-acid cycling drives nitrogen fixation in the legume–Rhizobium symbiosis. Nature. 422, 722– 726 (2003).

Maciejauskaite M. and Miliauskaite J. The efficiency of machine learning algorithms in classifying non-functional requirements. New Trends Comp. Sci. 2, 46–56 (2024).

Maged, M., Abdul A., and Heribert H. Tailoring plant-associated microbial inoculants in agriculture: a roadmap for successful application. J. Exp. Bot. 71, 3878–3901 (2020).

Martignoni, M., Garnier, J., Zhang, X., Rosa, D., Kokkoris, V., Tyson R. et al. Co-inoculation with arbuscular mycorrhizal fungi differing in carbon sink strength induces a synergistic effect in plant growth. J. Theor. Biol. 531, 110859 (2021).

McNew, G. The nature, origin, and evolution of parasitism. In Plant Pathology: An Advanced Treatise (eds Horsfall, J. G. & Dimond, A. E.) 19–69 (Academic Press, New York, 1960).

Milanese, A., Mende, D., Paoli, L., Salazar, G., Ruscheweyh, H., Cuenca, M. et al. Microbial abundance, activity and population genomic profiling with mOTUs2. Nat. Commun. 10, 1014 (2019).

Philippot, L., Chenu, C., Kappler, A., Rillig, M., Fierer N. The interplay between microbial communities and soil properties. Nat. Rev. Microbiol. 22, 226–239 (2024).

Poudel, R., Jumpponen, A., Schlatter, D., Paulitz, T., McSpadden Gardener, B., Kinkel, L. et al. Microbiome networks: a systems framework for identifying candidate microbial assemblages for disease management. Phytopathol. 106, 1083–1096 (2016).

Pozo, M., Cordier, C., Dumas-Gaudot, E., Gianinazzi, S., Barea, J., Azcón-Aguilar, C. Localized versus systemic effect of arbuscular mycorrhizal fungi on defense responses to Phytophthora infection in tomato plants. J. Exp. Bot. 53, 525–534 (2002).

R Core Team. R: A language and environment for statistical computing. R Foundation for Statistical Computing, Vienna, Austria. 2023. URL https://www.R-project.org/.

Raffa, K., Brockerhoff, E., Grégoire J., Hamelin, R., Liebhold A., Saninti, A. et al. Approaches to forecasting damage by invasive forest insects and pathogens: a cross-assessment. BioScience. 73, 85–111 (2023).

Rahman, S., Singh, E., Pieterse, C., Schenk, P. Emerging microbial biocontrol strategies for plant pathogens. Plant Sci. 267, 102–111 (2018).

Rheault, K., Lachance, D., Morency M., Thiffault, E., Guittonny M., Isabel, N. et al. Plant genotype influences physiochemical properties of substrate as well as bacterial and fungal assemblages in the rhizosphere of balsam poplar. Front. Microbiol. 11, 575625 (2020).

Rispail, N., and Rubiales, D. Identification of sources of quantitative resistance to *Fusarium oxysporum* f. Sp. Medicaginis in Medicago truncatula. Plant Dis. 98, 667–673 (2014).

Roorkiwal, M., Bharadwaj, C., Barmukh, R., Dixit, G., Thudi, M., Gaur, P. et al. Integrating genomics for chickpea improvement: achievements and opportunities. Theor. Appl. Genet. 133, 1703–1720 (2020).

Sasse J., Martinoia E., and Northen, T. Feed your friends: do plant exudates shape the root microbiome? Trends Plant Sci. 23, 25–41 (2018).

Sassone-Corsi, M. No vacancy: How beneficial microbes cooperate with immunity to provide colonization resistance to pathogens. J Immunol. 194, 4081–4087 (2015).

Savary, S., Willocquet L., Pethybridge, S., Esker, P., McRoberts, N., Nelson, A. The global burden of pathogens and pests on major food crops. *Nat*. Ecol. Evol. 3, 430–439 (2019).

Simonsen, A. K., and Stinchcombe, J. R. Standing genetic variation in host preference for mutualist microbial symbionts. Proc. R. Soc. B. 281, 20142036 (2014).

Singh, B., Delgado-Baquerizo, M., Egidi, E., Guirado, E., Leach, J., Liu, H. et al. Climate change impacts on plant pathogens, food security and paths forward. Nat. Rev. Microbiol. 21, 640–656 (2023).

Stone M. Cross-validatory choice and assessment of statistical predictions. J. Royal Stat. Soc. 36, 111–147, (1974).

Strange, R. and Scott, P. Plant disease: a threat to global food security. Annu. Rev. Phytopathol. 43, 83–116 (2005).

Subedi, S., Allen, P., Vidales, R., Sternberg, L., Ross, M., Afkhami, M. Salinity legacy: foliar microbiome’s history affects mutualist-conferred salinity tolerance. Ecology. 103, e3679 (2022).

Sultana, T., Deeba, F., Naz, F., Rose, R., Naqvi, S. Expression of a rice GLP in *Medicago truncatula* exerting pleiotropic effects on resistance against *Fusarium oxysporum* through enhancing FeSOD-like activity. Acta Physiol. Plant. 38, 225 (2016).

Sundin, G., Castiblanco, L., Yuan, X., Zeng, Q., Yang, C. Bacterial disease management: challenges, experience, innovation and future prospects: challenges in bacterial molecular plant pathology. Mol. Plant Pathol. 17, 1506–1518 (2016).

Tian, H., Kah, M., and Kariman, K. Are nanoparticles a threat to mycorrhizal and rhizobial symbioses? A critical review. Front. Microbiol. 10, 1660 (2019).

Trivedi, P., Leach, J., Tringe, S., Sa, T., Singh, B. Plant–microbiome interactions: from community assembly to plant health. Nat. Rev. Microbiol. 18, 607–621 (2020).

Tsuzuki, S., Handa, Y., Takeda, N. Kawaguchi, M. Strigolactone-induced putative secreted protein 1 is required for the establishment of symbiosis by the arbuscular mycorrhizal fungus *Rhizophagus irregularis*. MPMI. 29, 10.1094 (2016).

Tyerman, S., Niemietz, M., Bramley, H. Plant aquaporins: multifunctional water and solute channels with expanding roles. Plant Cell Environ. 25, 173 – 194 (2002).

Vigo, C., Norman, J., and Hooker, J. Biocontrol of the pathogen Phytophthora parasitica by arbuscular mycorrhizal fungi is a consequence of effects on infection loci. Plant Pathol. 49, 509–514 (2000).

Waller, L., Smith, D., Childs, J., Real, L. Monte Carlo assessments of goodness-of-fit for ecological simulation models. Ecol. Modell. 164, 49–63 (2003).

Wang, R., Wang, M., Chen, K., Wang, S., Alejandro Jose Mur, L., Guo S. Exploring the roles of aquaporins in plant-microbe interactions. Cells. 7, 267 (2018).

Williams A., Sharma, M., Tatcher, L., Azam, S., Hane, J., Sperschneider, J., et al. Comparative genomics and prediction of conditionally dispensable sequences in legume-infecting *Fusarium oxysporum* formae speciales facilitates identification of candidate effectors. BMC Genom. 17, 191 (2016).

Young, N and Udvardi, M. Translating *Medicago truncatula* genomics to crop legumes. Curr. Opin. Plant Biol. 12, 193–201 (2009).

Zhong, Y., Yang, Y., Liu, P., Xu, R., Rensing, C., Fu, X. et al. Genotype and rhizobium inoculation modulate the assembly of soybean rhizobacterial communities. Plant Cell Environ. 42, 2028–2044 (2019).

Zubrod, J., Bundschuh M., Arts, G., Brühl, C., Imfeld, G., Knäbel, A. et al. Fungicides: an overlooked pesticide class? ES&T. 53, 3347–3365 (2019).

