## supplementary_figures for "Non-additive interactions between multiple mutualists and host plant genotype simultaneously promote increased plant growth and pathogen defense"

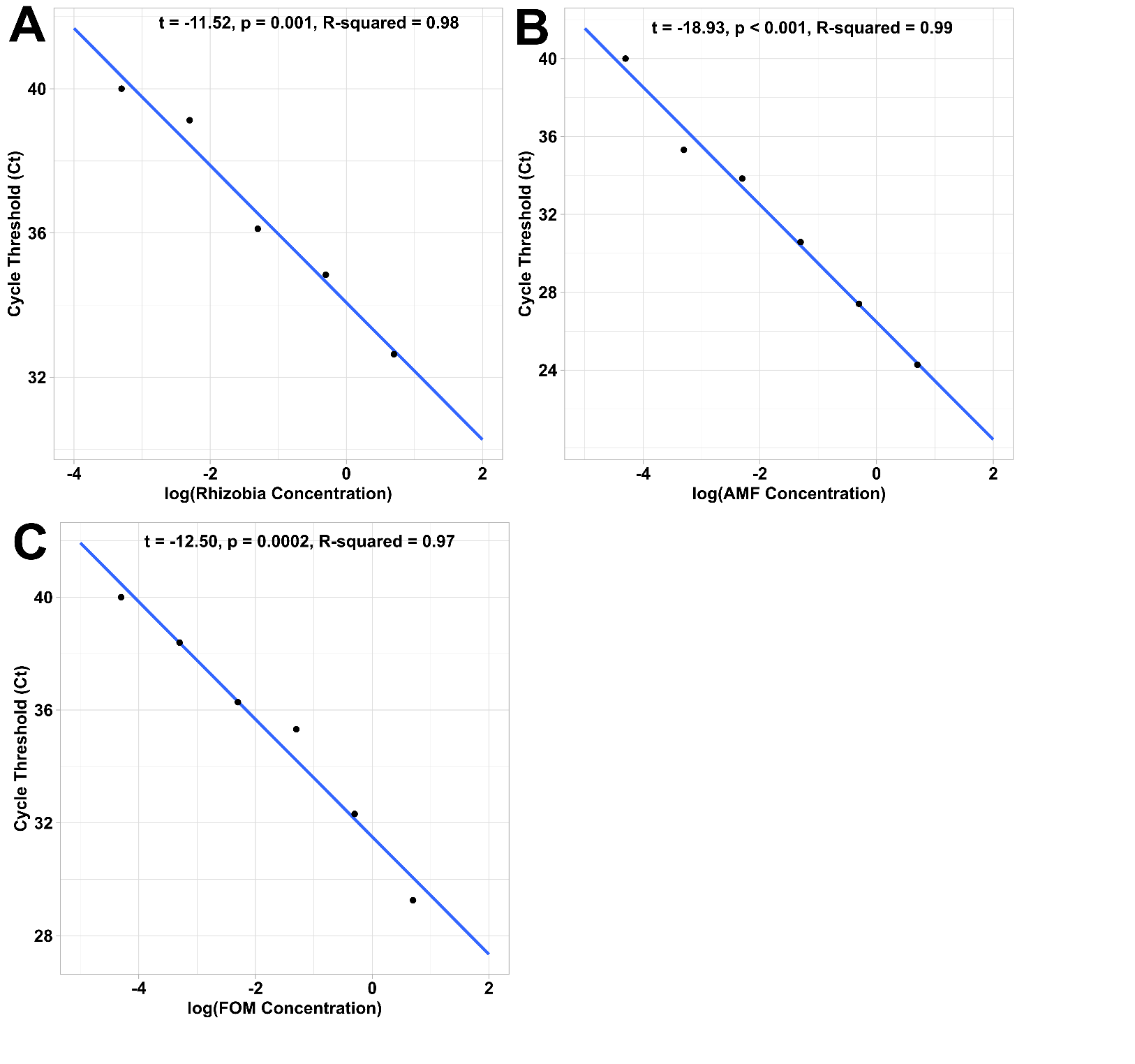
**Fig. S1. Standard dilution curves for microbial abundances.** Standard dilution curves for estimation of microbial abundances for **(A)** rhizobia, **(B)** AMF, and **(C)** FOM based on qPCR cycle threshold data. Concentration is displayed with log10 units.

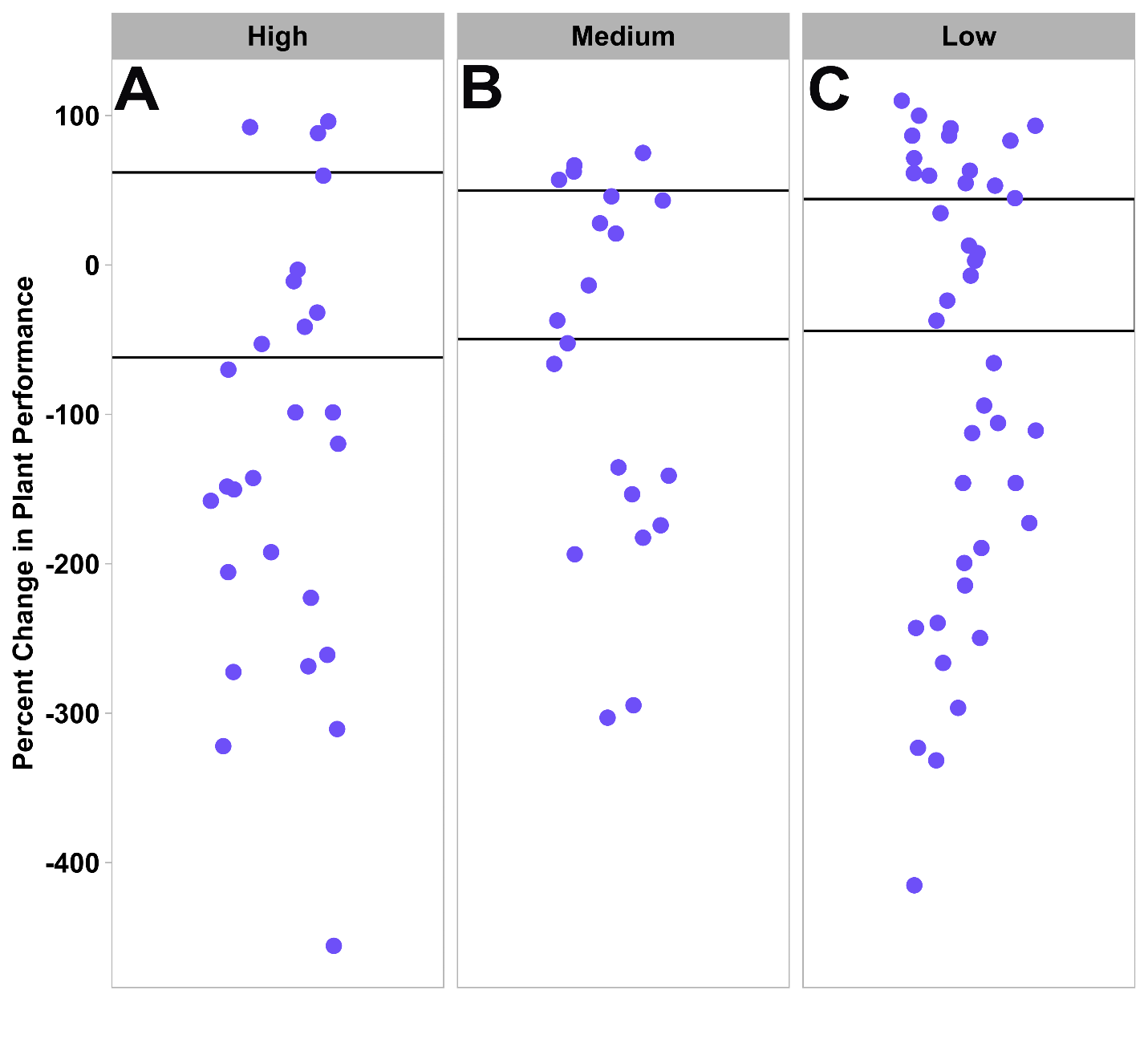

**Fig. S2. Most individuals showing an antagonistic response to multiple mutualists performed the same or worse than controls.** Percent change in plant performance of plants showing an antagonistic response to multiple mutualists for genotypes classified as having **(A)** high, **(B)** medium, or **(C)** low tolerance to general fungal pathogen invasion. Each point represents the change in plant performance of a biological replicate with antagonism inoculated with multiple mutualists (R+M+) relative to controls with no mutualists (R-M-) measured in a benign environment at 5 weeks post inoculation with mutualists. Percent change in plant performance axis values are shared across all panels. The bold lines demarcate the 95% CI of the control group. Points that fall above the boundary performed better than the controls, points that are within the boundary performed the same as controls, and points that are below the boundary performed worse than controls.

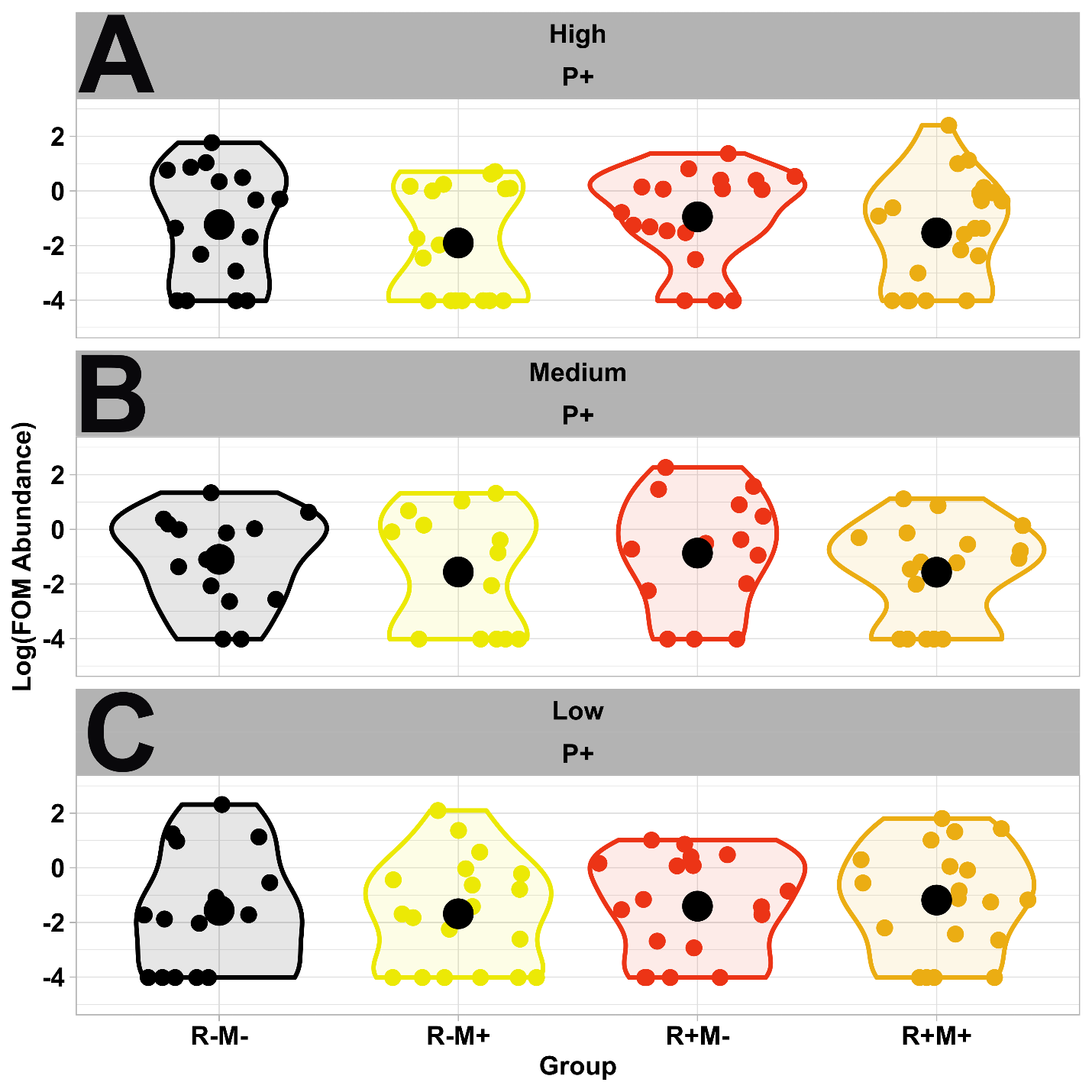

**Fig. S3. Fusarium abundance was the same across all mutualist treatments and tolerance types.** Violin plot where each point represents the log fusarium abundance of a biological replicate and large black circle denotes the mean. Panels split the data by tolerance cluster (i.e. high, medium, and low) and mutualist treatment where the color indicates the mutualist inoculation status; yellow are plants with AMF being the only mutualist, red are plants with rhizobia being the only mutualist, orange are plants with both AMF and rhizobia, and black are controls with no mutualists.

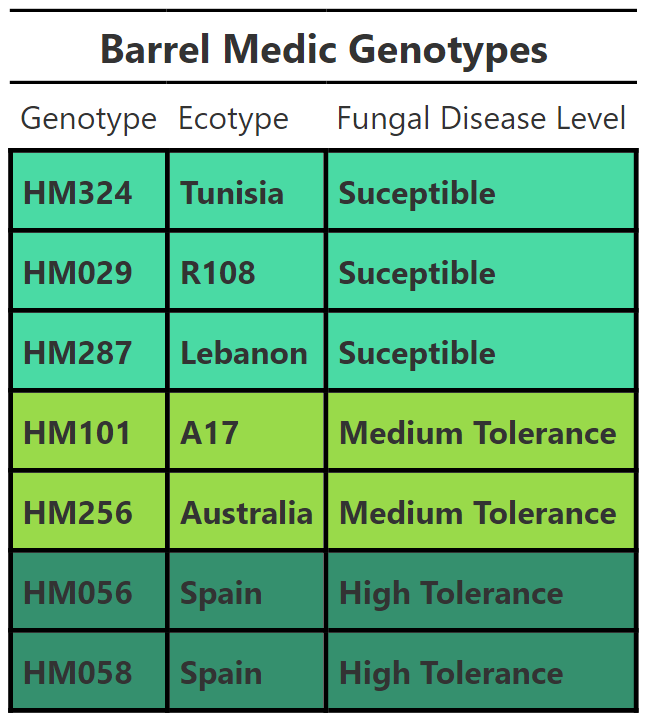

**Table S1. 7 unique *M. truncatula* genotypes with differing levels of tolerance against general fungal disease used in this study.** Fungal disease tolerance ranged from susceptible (highlighted in red), medium tolerance (highlighted in orange), and high tolerance (highlighted in yellow).

| **Variables** | **Dfn** | **Dfd** | **F** | ***p*** |
| --- | --- | --- | --- | --- |
| **Plant Performance – High Tolerance** |  |  |  |  |
| Timepoint | 1 | 85 | 16.21 | **0.0001** |
| Environment | 1 | 85 | 0.03 | 0.87 |
| Timepoint:Environment | 1 | 85 | 0.11 | 0.75 |
| **Plant Performance – Medium Tolerance** |  |  |  |  |
| Timepoint | 1 | 57 | 0.89 | 0.35 |
| Environment | 1 | 57 | 0.03 | 0.86 |
| Timepoint:Environment | 1 | 57 | 0.03 | 0.87 |
| **Plant Performance – Low Tolerance** |  |  |  |  |
| Timepoint | 1 | 73 | 7.47 | **0.008** |
| Environment | 1 | 73 | 5.16 | **0.03** |
| Timepoint:Environment | 1 | 73 | 3.92 | **0.05** |
| **Plant Performance – Low Tolerance - Benign** |  |  |  |  |
| Timepoint | 1 | 38 | 8.19 | **0.007** |
| **Plant Performance – Low Tolerance - Pathogenic** |  |  |  |  |
| Timepoint | 1 | 35 | 0.50 | 0.49 |

**Table S2. Factors affecting plant performance.** Repeated measure ANOVAs were used to assess plant performance (composite metric for plant growth) for significant effects of timepoint, environment, and their interaction. Significant *p*-values are highlighted in bold. Since the interaction term was significant for the low tolerance group additional ANOVAS were broken into simple effects by environment type.
